# A Mechanistic Understanding of the Activity-Stability Trade-off in PETase

**DOI:** 10.1101/2024.06.09.598049

**Authors:** Shuang Chen, Ekram Akram, Weili Qiao, Yifei Zhang, Shozeb Haider, Yufei Cao

## Abstract

Enzymatic degradation of poly(ethylene terephthalate) (PET) has garnered considerable interest in plastic recycling efforts. Despite numerous descriptions of both natural and engineered enzymes, the fundamental mechanism underlying PETase-catalyzed PET depolymerization at the solid-liquid interface remains elusive. This lack of understanding hampers the rational design of highly efficient depolymerases. Here, we employ multiscale simulations and experiments to elucidate the complete catalytic pathway of *Is*PETase, from enzyme adsorption at the interface to PET fragment capture, conformational refinement, and ester bond cleavage. Both endo- and exo-cleavage modes of the enzyme are identified, indicating its capacity for endo- and exo-lytic activities. We discover that the trade-off between the activity and stability of *Is*PETase’s PET-capturing pliers brings compromises to its PET depolymerization performance. Reshaping the loop dynamics of the enzyme can break this trade-off and enhance its stability and activity simultaneously, as demonstrated by the evolved variant HotPETase. Overall, our study offers comprehensive details into how PETase functions at the interface and provides valuable insights for engineering efficient plastic-degrading enzymes.

## Introduction

The global environmental crisis caused by plastic waste has driven extensive research into finding sustainable solutions for plastic recycling and upcycling^1-4^. Among these efforts, the enzyme-catalyzed deconstruction of poly(ethylene terephthalate) (PET) has emerged as a promising approach^5^, due to its mild pH and relatively lower temperature conditions for recycling PET into monomers. Numerous natural and engineered enzymes, known as PET hydrolase or PETase, have been reported for PET biodegradation^4,6-10^.

PETase specifically targets the ester bonds of the PET chain, facilitating monomer recovery. Unlike most enzymes that function in homogeneous aqueous environments, PETase operates at the solid-liquid interface, presenting a significant challenge for PETase engineering and mechanistic studies. Most strategies for enhancing PETase efficiency and elucidating its catalytic mechanism rely on unrealistic substrate fragments^4,11-14^. Despite extensive experimental and theoretical research, the mechanism underlying PETase-catalyzed PET depolymerization at the solid-liquid interface, including polymeric substrate accessibility, enzyme-substrate complexation, its endolytic or exolytic activity, and enzyme stability, remains poorly understood^5^. The detailed interactions between macromolecular PET substrates and enzyme have yet to be reported, posing a significant barrier to the rational design of highly efficient engineered PETase.

Prior to the discovery of PETase, a class of enzymes called cellulases that also hydrolyze insoluble polymeric substrates had been extensively studied^15^. For instance, exocellulases target the crystalline regions of cellulose fibers, exhibit processive catalysis, and are directionally dependent when cleaving two to four units from the ends of exposed chains^16^. A cellulase typically consists of a carbohydrate-binding module (CBM), a larger catalytic domain, and a linker domain connecting the two^17,18^. Unlike cellulases, natural PETase lacks a polymer-binding module and primarily degrades the amorphous region of PET^9^. Consequently, PETase does not function in a processive and directional manner like cellulase. Furthermore, grafting cellulase’s CBM onto PETase has been shown to have minimal impact on its efficiency^19^. The mode of PET degradation by PETase, whether endo- or exo-type, remains debatable. Mass spectrometry analysis suggests endo-type hydrolysis^20,21^, while NMR spectroscopy indicates both endo- and exo-type activities^22^. A recent study by Bell et al. proposed an exo-cleavage mechanism, supported by minimal changes in the molecular weight and dispersity of residual PET^9^.

To reveal the mechanism of PET biodegradation, we established a realistic PET-water interface model to investigate enzyme adsorption, conformational changes, and ester bond recognition using *Idonella sakaiensis* PETase (*Is*PETase^6^) as a model enzyme. Our findings revealed that *Is*PETase exhibits non-specific binding to the PET surface, which could lead to enzyme inactivation due to the inherent instability of the catalytic loop, as confirmed by experimental evidence. Through extensive sampling, we identified both endo- and exo-cleavage conformations of enzymes and elucidated the detailed processes of ester bond recognition. By comparing *Is*PETase with the evolved thermostable variant HotPETase, we demonstrated how directed evolution breaks the activity-stability trade-off and enhances the PET depolymerization performance of the enzyme. This trade-off appears to be universal in PETases, as evidenced by studies on engineered leaf-branch compost cutinase (LCC), another extensively investigated PETase. Our work provides guiding principles for modulating loop dynamics in order to create robust plastic-degrading enzymes.

## Results and Discussion

### PET substrate recognition by *Is*PETase

We began by constructing a solid-liquid interface model to investigate the mechanism, incorporating an amorphous PET membrane and water (Supplementary Fig. 1-5). An unbiased protocol of supervised molecular dynamics (MD) simulations was developed to sample the binding states of enzymes on the PET surface (see details in the Methods section). During supervised MD, the enzyme gradually approached and eventually bound to the PET surface (Supplementary Fig. 6). After over 100 repeated samplings, we examined the distribution of enzyme adsorption states on the PET surface (Fig. 1a). The *Is*PETase adsorption did not display obvious specificity, as in most cases, the active sites of enzymes were not oriented towards the PET surface. The binding free energy (ΔG_binding_) distribution illustrates that the average ΔG_binding_ values of active states were slightly higher than those of the non-active states, further confirming the limited targeted binding capacity of *Is*PETase (Fig. 1b-c and Supplementary Fig. 7-8).

**Figure 1.**
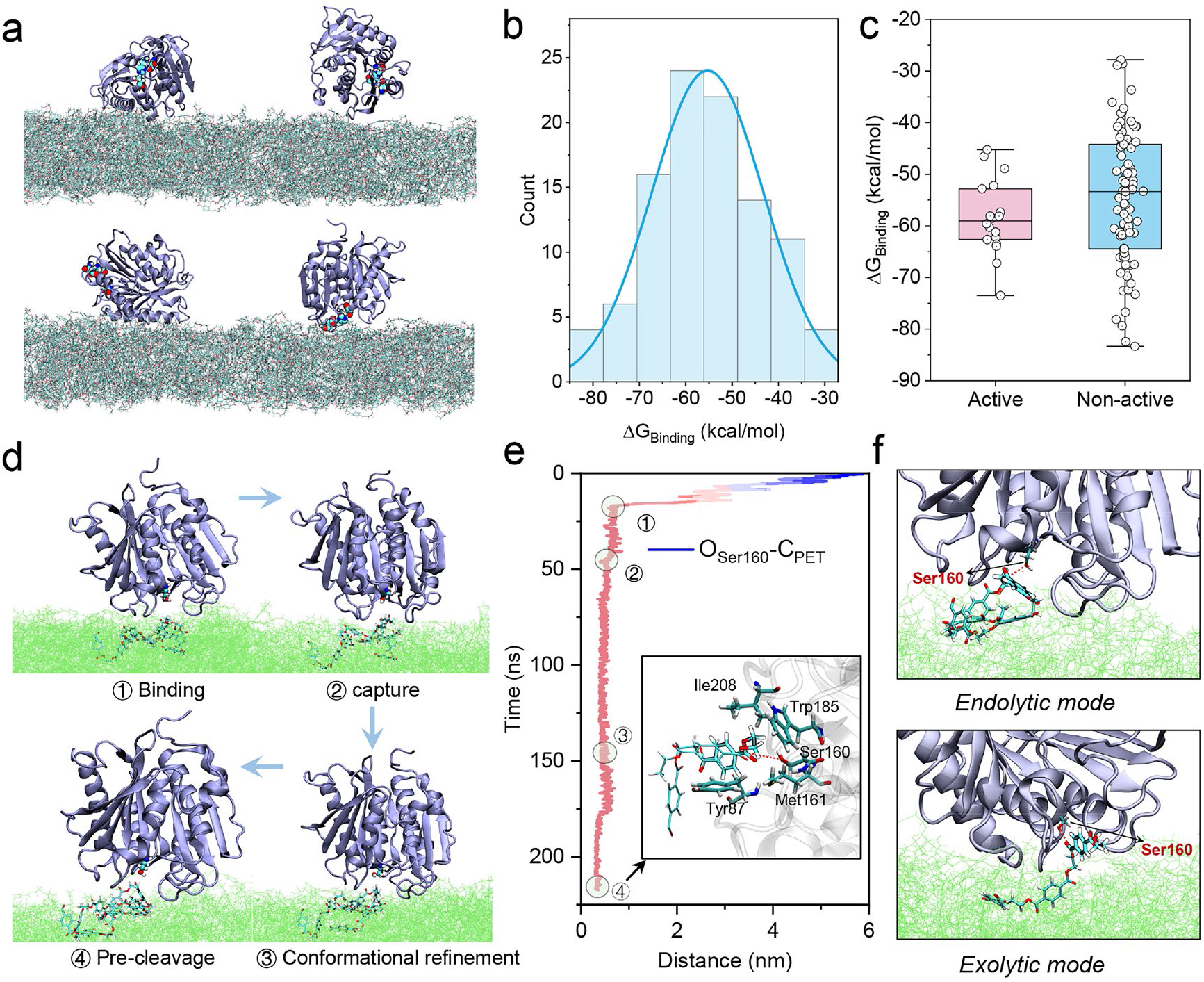
PET recognition by *Is*PETase at a solid-water interface. (a) Various binding states of *Is*PETase on the PET surface, with the catalytic triad shown in a ball representation. (b) Distribution of binding free energy across all binding states. (c) Binding free energy of active and non-active binding states. Binding was defined as “active” if the distance between Ser160 and the PET surface was less than 5.5 Å in the final binding states. (d) Detailed procedure of ester bond recognition by *Is*PETase. Ser160 and the captured PET fragments are shown in ball and stick representations, respectively. (e) Evolution of the distance between the side chain hydroxyl of Ser160 and the carbon atom of the recognized PET ester bond. The final pre-cleavage conformation was illustrated. (f) Endolytic and exolytic pre-cleavage modes of *Is*PETase.

The simulations depicted the detailed mechanism of ester bond recognition by the enzyme (Fig. 1d-e, Supplementary Fig. 9-11), which involves four steps: enzyme binding, PET fragment capture, conformational refinement, and attainment of the final pre-cleavage conformation. Throughout this process, the distance between Ser160 in the active site and the ester bond of PET gradually decreased. Residues within the loops near the active site, including Tyr87, Leu117, Trp159, Met161, Trp185, and Ile208 played a crucial role in capturing the PET fragments (Supplementary Fig. 11). This was consistent with experimental results (Supplementary Fig. 11d). Notably, Tyr87, Trp185 and Ile208, located on two loops, acted as “pliers” for capturing PET (Fig. 1e). We identified two distinct PET-capturing states of *Is*PETase, the endolytic and exolytic modes, indicating that *Is*PETase possesses both endolytic and exolytic activities (Fig. 1f, Supplementary Fig. 12-13). This suggests that both two catalytic modes function cooperatively during PET depolymerization.

### The trade-off between capturing flexibility and stability

Upon monitoring the catalytic triad conformation of *Is*PETase during PET recognition, we observed three types of conformational dynamics: stable, fluctuated, and destroyed (Fig. 2a-b, Supplementary Fig. 14-15). Thus, PET binding could perturb the conformation of *Is*PETase and even lead to enzyme inactivation. The disrupted states of the enzyme can potentially be restored or permanently deactivated (Supplementary Fig. 16). To investigate the origin of PET-induced *Is*PETase inactivation, we conducted a 1 μs-long MD simulation of the enzyme’s unbound state. It revealed the inherent instability of *Is*PETase, with significant fluctuations observed in the conformation of the enzyme’s active site (Fig. 2c). As shown by the root mean square fluctuations (RMSF), the catalytic loop containing Asp206 exhibited high flexibility (Fig. 2d-e). This flexibility, which could lead to a loss of active site preorganization, is the possible cause of *Is*PETase’s inherent instability. We experimentally investigated the deactivation of *Is*PETase in the presence of PET powders using *p*-nitrophenyl butyrate (*p*NPB) as the substrate. The results indicated that the adsorption of PETase on either low-crystallinity PET (LC-PET) or high-crystallinity PET (HC-PET) accelerates the deactivation of the enzyme, consistent with the MD simulations (Fig. 2f).

**Figure 2.**
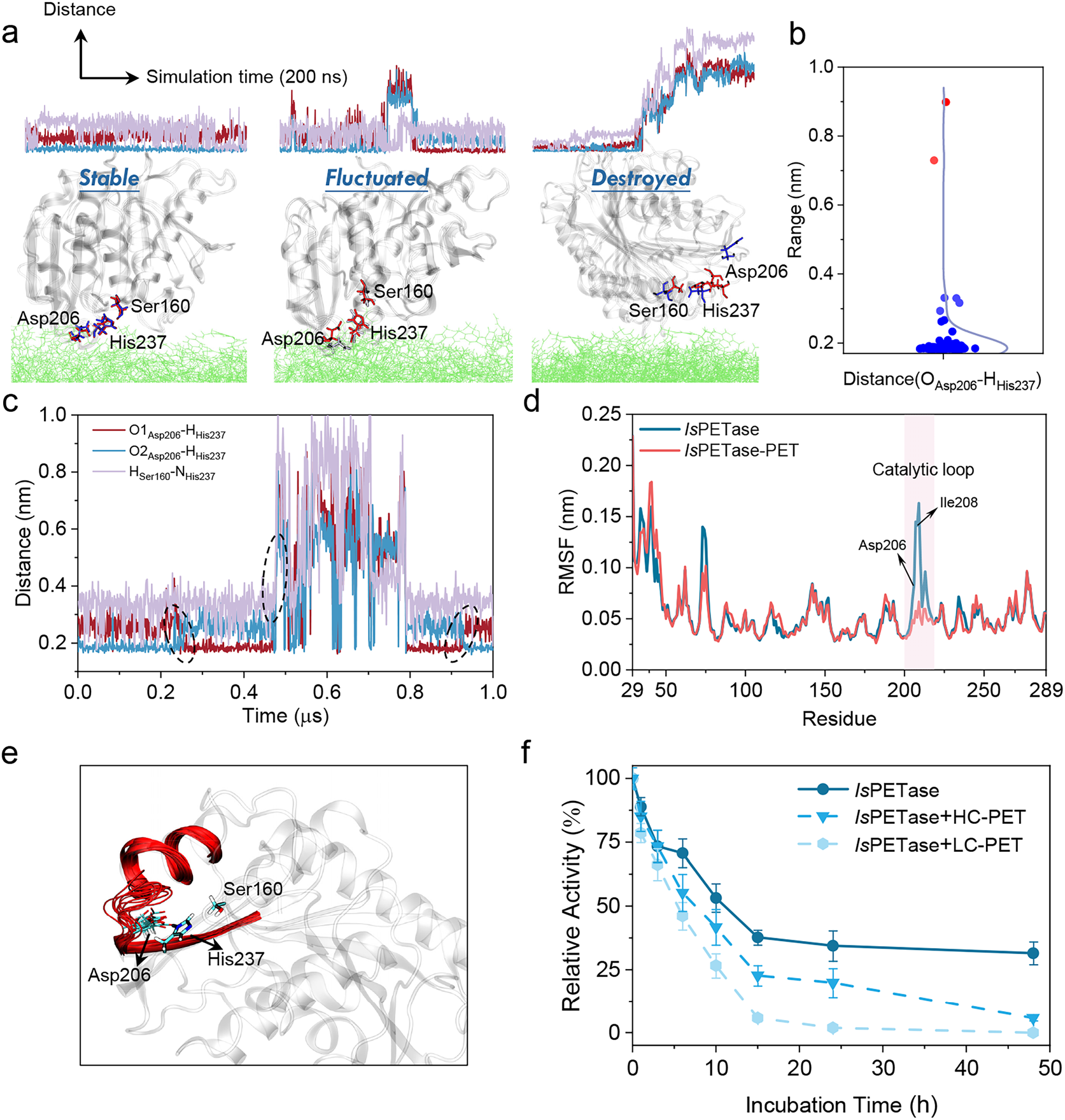
Inherent instability of *Is*PETase. (a) Three types of conformational dynamics of the enzyme active site in PET-bound states. (b) Distribution of the distance between the oxygen atom of Asp206 and the hydrogen atom of His237. (c) Evolution of three pairs of distances within the active site of the unbound enzyme. (d) RMSF of unbound and PET-bound *Is*PETase. (e) Conformational fluctuations of the enzyme catalytic loop. (f) Time-course deactivation of *Is*PETase when adsorbing on LC-PET and HC-PET powders.

In the PET-bound states, the flexible loop and Trp185, which form one side of the PET capture pliers, adopted a confined conformation. They tightly interacted with the PET fragments (Fig. 2d, Supplementary Fig. 17-21). We speculated that the flexibility of the catalytic loop and the wobbling of Trp185 contribute to the PET capture capability of *Is*PETase, despite inducing instability. Previous studies by Guo and colleagues showed that the introduction of Trp185 wobbling in *Is*PETase can enhance the activity of other PETases, albeit at the cost of inducing instability^23^ (Supplementary Note 1, Supplementary Fig. 22-23). Taken together, there exists a trade-off between PET capture capacity and the stability of *Is*PETase. This trade-off may explain *Is*PETase’s outstanding PET depolymerization activity at moderate temperatures and its instability at higher temperatures^6^.

### Directed evolution breaks the trade-off

Recent work by Bell et al. created a thermostable variant of *Is*PETase known as HotPETase^9^. HotPETase incorporates 21 mutations, including three from the original protein template, two from the rational insertion of an additional disulfide bridge, and 16 identified through directed evolution (Fig. 3a). Analysis of the ΔG_binding_ distribution revealed that HotPETase exhibited a slightly higher binding capability than the wild type enzyme (Fig. 3b, Supplementary Fig. 24-25), with more states characterized by high ΔG_binding_ value. Adhesion force measured by atomic force microscopy-based single-molecule force spectroscopy (AFM-SMFS) showed a consistent trend of binding affinity of *Is*PETase and HotPETase to a PET film (Fig. 3c). The difference in ΔG_binding_ between active and non-active states closely resembled that of *Is*PETase (Supplementary Fig. 25), indicating that directed evolution did not lead to an improvement in the target binding ability. Notably, all binding states of HotPETase adopted a stable conformation of the catalytic triad, underscoring the significantly improved stability of the enzyme (Fig. 3d, Supplementary Fig. 26-27). Microsecond-long MD simulation and conformational landscape analysis of the unbound enzyme further confirmed the intrinsic stability of HotPETase (Supplementary Fig. 28). Our experiments also showed that HotPETase exhibits greater stability than *Is*PETase, both in its free form and in the presence of PET powder (Fig. 3e, Supplementary Fig. 29). As shown in Fig. 3f, the fluctuation of the catalytic loop was greatly reduced. This reduction was attributed to the salt bridge and hydrogen bond interactions between Arg207 and its neighboring residues (Glu204, Asn205, and Asp206), as well as steric hindrance from Tyr214 (Fig. 3g, Supplementary Fig. 30). It is noteworthy that Arg207 and Tyr214 are two of the three mutations in the initial round of directed evolution for HotPETase^9^, the round resulting in the largest enhancement of T_m_ (Supplementary Fig. 31). Thus, both experimental and simulation results unanimously affirm the pivotal role of Arg207 and Tyr214 in enhancing the enzyme’s stability.

**Figure 3.**
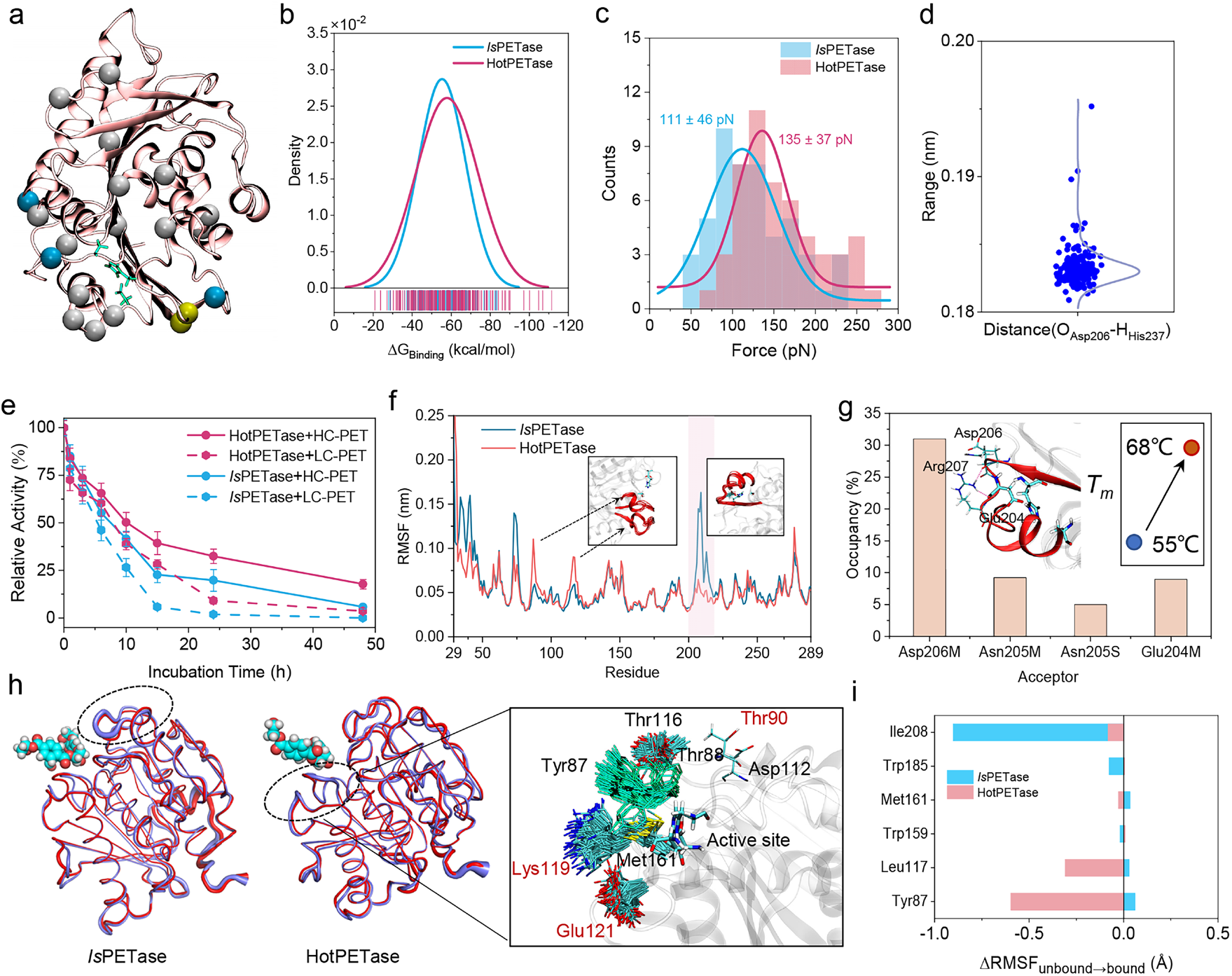
Intrinsic Stability of HotPETase. (a) Comparison of mutations in HotPETase with *Is*PETase: three positions in the starting template (blue spheres), a rationally installed disulfide bridge (yellow spheres), and 16 positions identified through evolution (white spheres). The catalytic triad was depicted with a stick representation in green. (b) Distribution of binding free energy for all binding states of *Is*PETase and HotPETase. (c) Histograms of detachment force distribution of *Is*PETase and HotPETase interacting with the surface of a PET film. The mean adhesion force was obtained by a Gaussian fitting. (d) Distribution of distances between the oxygen atom of Asp206 and the hydrogen atom of His237. (e) Time-course deactivation of *Is*PETase and HotPETase when adsorbing on LC-PET and HC-PET powders. (f) RMSF analysis of *Is*PETase and HotPETase. (g) Hydrogen bonds induced by mutations located in the catalytic loop. The donor of hydrogen bonds are Arg207 and “M” and “S” denote the mainchain and sidechain of residues. (h) Two distinct wobbling modes observed in *Is*PETase and HotPETase. The unbound and bound enzymes were shown in blue and red, respectively. (i) The ΔRMSF of several key residues from unbound state to bound state.

The current simulations and previous analysis of the HotPETase crystal structure both reveal a missing Trp185 wobbling in HotPETase (Supplementary Fig. 32) and low flexibility of the catalytic loop (Fig. 3f). Hence, the flexibility of one jaw of the PET-capturing pliers containing Trp185 and Ile208 was significantly reduced. However, we discovered that Tyr87 on the other jaw of the pliers and its associated loop exhibited higher flexibility compared to the wild type, indicating a new wobbling residue (Fig. 3f, g-h, Supplementary Fig. 33-35). This originates from three mutations at Thr90, Lys119 and Glu121, leading to a broken salt bridge and less steric hindrance (Supplementary Fig. 36-38). Directed evolution compromised the flexibility of one side of the PET-capturing pliers to improve enzyme stability, while simultaneously introducing wobbling in Tyr87 and enhancing the flexibility of the opposite side of the pliers to offset this loss. Consequently, the reshaped conformational landscape of HotPETase overcomes the limitation of the activity-stability trade-off associated with the flexibility of the catalytic loop and Trp185 wobbling. We propose that Tyr87 wobbling could enhance the activity of other PETases and exhibit greater robustness at elevated temperatures.

### Origin of HotPETase’s high intrinsic activity and PETase designing opportunities

In most experiments involving PET degradation by PETase, enzyme activity is assessed based on the accumulation of mono(2-hydroxyethyl) terephthalate (MHET) and terephthalic acid (TPA) over several hours. Our simulations of the catalytic process of PETase reveal that the apparent enzyme activity reflects not only intrinsic depolymerization activity, but also PET capture capability and stability. Thus, the enhanced catalytic performance cannot be attributed to a single factor. This is supported by the observed improvement in PET depolymerization performance and modest changes (even decrements) in activity towards soluble small substrates in engineered PETase^24^.

To elucidate the contribution of directed evolution, we compared the intrinsic activity between the wild-type *Is*PETase and its evolved variant HotPETase. QM/MM calculations of PET acylation, the rate-limiting step in *Is*PETase catalysis^25,26^, demonstrated a lower free energy barrier for HotPETase-catalyzed PET trimer hydrolysis compared to *Is*PETase (Fig. 4a-b, Supplementary Fig. 39). The heightened activity of HotPETase stems from a favorable enthalpic contribution and lower entropic cost (Fig. 4c and Supplementary Table 1). Analysis of the conformational dynamics of PET-capturing pliers (distance between two loops) indicated that HotPETase adopts a more fluctuant conformation at ground states (GS) and a more confined conformation at transition states (TS) than *Is*PETase (Fig. 4d-e, Supplementary Fig. 40). This suggests a more effective TS stabilization by the exquisite active-site preorganization of HotPETase (Supplementary Fig. 41). Hence, the transition from a flexible GS to a rigid TS could lead to activity gains.

**Figure 4.**
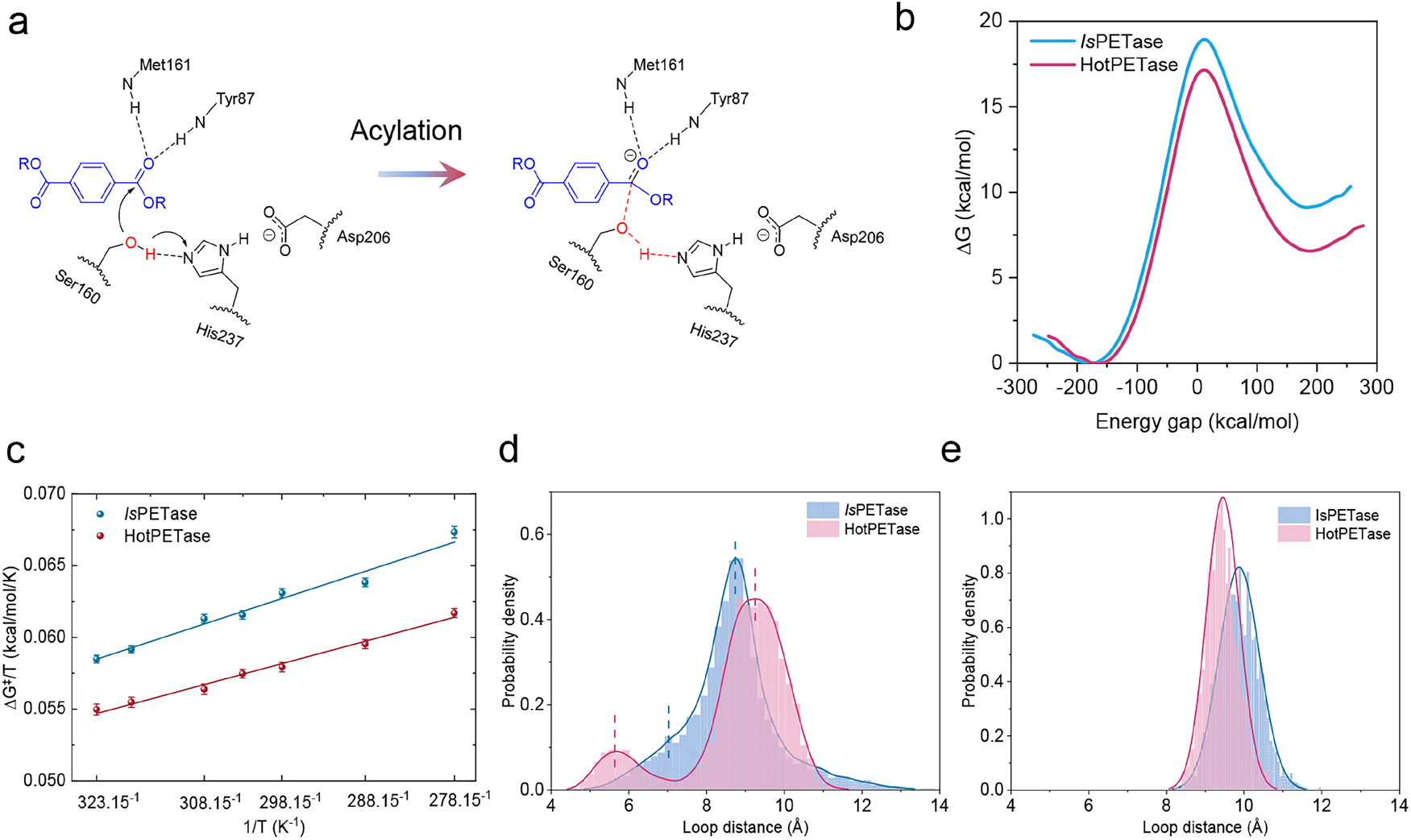
High intrinsic activity of HotPETase. (a) Acylation step of *Is*PETase and HotPETase. (b) Calculated free energy profiles (298.15 K) from reactants to the tetrahedral intermediate in the acylation step. (c) Arrhenius plots depicting ΔG^‡^/T vs. 1/T for catalyzed reactions in *Is*PETase and HotPETase. (d) The loop distance between residues Tyr87 (C_*γ*_ atom) and Ile208 (C_*β*_ atom) at (d) free enzyme states and (e) enzyme-substrate transition states.

Drawing insights from directed evolution, tuning the conformational dynamics of PET-capturing pliers of PETase emerges as an effective approach for enzyme engineering. The intrinsic activity, PET capture capability, and enzyme stability are influenced by the dynamics of abundant loops near the enzyme’s active site. Comparing the differences in loop dynamics between different PETases and identifying the key residues that modulate them will facilitate the rational design of highly active and stable PETase enzymes.

## Conclusion

Our multiscale MD simulations and experiments demonstrate how PETase functions at a solid-liquid interface. The dynamic loops near the enzyme active site play a crucial role in capturing polymer substrate fragments. However, the activity-stability trade-off poses a challenge to enhancing enzyme performance. The catalytically superior HotPETase overcomes this trade-off by altering loop dynamics through directed evolution. By shifting flexibility from one side of the PET-capturing pliers to the other, it achieves improvements in both enzyme activity and stability. The comprehensive understanding of PETase’s catalytic process provided by our simulations will contribute to elucidating the mechanism of enzyme catalysis at interfaces and to engineering robust plastic-degrading enzymes.

## Methods

### Building the PET membrane model

Parameters for the PET monomer were generated with the antechamber module of Amber18^27^ using the general Amber force field (GAFF)^28^. The partial charges were set to fit the electrostatic potential generated with B3LYP/TZVP by RESP^29^. Details of the repeat units and tail end of the polymer chain can be found in Supplementary Fig. 1. Initially, twelve PET chains (PET_100_) were placed in a large periodic cubic box and simulated under vacuum for 50 ns using the NVT ensemble with a V-rescale thermostat^30^ set at 800 K. Once a complex structure of PET chains was obtained, pressure was added along the z-axis of the simulation box. The x and y directions of the simulation box had periodic boundary conditions, while z=0 and z=box positions were occupied by two walls composed of CA atoms (atom type in the Amber force field). Under pressure in the z direction, the PET complex was compressed into a membrane. To achieve homogeneity of different PET chains within the membrane, we conducted simulations for 100 ns at 800 K followed by an additional 100 ns simulation at 300 K. During the simulation, an anisotropic Berendsen barostat was used with a reference pressure of 1 bar. The solid PET membrane was then immersed in a water box, with water molecules represented by the three-point charge TIP3P model. Subsequently, the polymer-water system was equilibrated for 10 ns at a constant temperature of 298.15 K and pressure of 1 bar (NPT ensemble), initially by restraining the positions of polymers followed by NPT equilibration for another 200 ns without position restraints.

### MD simulation of the enzyme-PET complex

The crystallographic structure of the *Is*PETase (PDB ID: 5xjh)^11^ and HotPETase (PDB ID: 7qvh)^9^ were used as the starting coordinates. PROPKA 3^31,32^ was utilized to assign the protonation states of titratable residues, with further validation through visual inspection of protonation states and side chain orientations. The His residue of the catalytic triad was singly protonated at *N_σ_*. All crystal waters were removed. Enzyme molecules were immersed in a water box containing a PET membrane, as illustrated in Supplementary Fig. 6, with the TIP3P water model applied. All classic MD simulations were conducted using the GROMACS 2019.3^33^, along with the Amber ff14SB^34^ force-field parameters for protein. The entire system underwent energy minimization using a combination of steepest descent and conjugate gradient methods (maximum force < 100 kJ/mol/nm). Subsequently, the solvent was equilibrated for 20 ns at constant temperature (298.15 K) and pressure (1 bar) under the NPT ensemble, with enzyme positions restrained. The resulting equilibrated structure served as the initial state for subsequent supervised MD simulations. Temperature and pressure were maintained using a V-rescale thermostat and Parrinello−Rahman barostat^35^, respectively. The cutoff radius for neighbor searching and nonbonded interactions was taken to be 12 Å, and all bonds were constrained using the LINCS algorithm^36^. Visualization of enzyme structures was accomplished using VMD^37^.

### Supervised MD simulation

The simulation protocol was adapted from a previous study on ligand-receptor interactions^38^. During the production of the MD trajectory, the z-coordinate of the enzyme mass center was monitored over a fixed time window (1 ns). A linear function *z*(*t*) = a\**t* +b was fitted to the z-coordinates as a function of simulation time. If the slope (a) was less than zero, indicating a decrease in the enzyme-PET distance during the simulation window, the MD simulation was restarted from the last set of coordinates. Otherwise, the simulation was resumed from the original set of coordinates and velocities, and the MD sampling was repeated. If the number of restarts exceeded five, the repeated sampling was terminated, and the MD simulation was restarted from the last set of coordinates. The supervision algorithm continued until three criteria were met: a negative slope, more than 80 contacts between the enzyme and PET (with a distance cutoff of 3.5 Å), and a z-coordinate of the enzyme mass center less than 65 Å. Following the supervised MD, an extended 200 ns-long NPT MD simulation was conducted to obtain the final equilibrated enzyme-PET complex.

### Analysis of MD simulation results

The binding free energy between the enzyme and PET surface, as well as energy decomposition, was assessed using the gmxMMPBSA^39^. Side-chain torsion angles and distances between different protein atoms were calculated using the Python library MDTraj^40^. We analyzed side-chain torsion angles up to χ_2_ and employed the Jensen-Shannon (JS) divergence to quantitatively compare the influence of PET binding on the probability distribution of side-chain torsion angles^41^. Using JS divergence, we visualized the change in the probability distribution of side-chain torsion angles of each residue after PET binding. When both χ_1_ and χ_2_ of a residue existed, the larger JS divergence was chosen. Additionally, time-lagged independent component analysis (tICA) was performed using PyEMMA 2^42^ to analyze the enzyme’s conformational landscape, with side-chain torsion angles of several key residues utilized as features (Supplementary Fig. 21).

### QM/MM calculations

Enzyme-substrate complexes were immersed within a spherical water droplet using the TIP3P water model^43^, with a radius of 42 Å. The choice of these radii ensured that all atoms of enzyme-substrate complexes resided within the inner 85% of the spherical boundary. All calculations utilized the OPLS-AA force field^44^. van der Waals (vdW) and bonding parameters for PET were derived from Schrödinger’s Macromodel, with partial charges assigned through the RESP protocol^45^. Water molecules close to the spherical boundary underwent radial and polarization restraints, in accordance with the SCAAS model^46^. All QM/MM (EVB/MD calculations) were conducted utilizing the Q6 program^47^, implementing the leapfrog integrator with a time step of 1 fs. Long-range interactions were managed employing the local reaction field (LRF) approach^48^ with a nonbonded interactions cutoff of 10 Å. A nonbonded interaction cutoff of 99 Å was applied specifically to the reacting fragments. The system temperature was maintained at a constant level through the utilization of the Berendsen thermostat^49^, employing a 100 fs bath coupling time. All simulation systems underwent a stepwise heating process from an initial temperature of 1 K to their respective final temperatures over a duration of 230 ps, subsequently followed by an equilibration period of 100 ps. The SHAKE algorithm^50^ was used to constrain bonds and angles of solvent molecules. EVB/MD free energy perturbation (FEP) simulations were conducted with 51 discrete λ windows, each with a duration of 10 ps. The simulations were initiated at λ1 = λ2 = 0.5, in proximity to the transition state, and progressed towards the reactant state (λ1 = 1) and the intermediate state (λ2 = 1). To prevent excessive dissociation of reacting fragments within the Michaelis and product complexes, a modest restraint of 0.1 kcal/mol/Å^2^ was applied to the reactive atoms throughout the EVB simulations. To calibrate the EVB reaction potential energy surface, the average free energy profile from 100 independent simulations for the reference reaction system in water was fitted to the corresponding free energy profile obtained by the DFT cluster calculations at 298.15 K. The temperature dependence of the reaction catalyzed by enzyme was analyzed by repeating the free energy profile calculations at 278.15, 288.15, 298.15, 303.15, 308.15, 318.15, and 323.15K. At each temperature, about 100 randomized individual replicate simulations were carried out to obtain the free energy profiles.

### Stability of enzymes

The stability of *Is*PETase and HotPETase in the soluble and adsorbed states was evaluated by their hydrolytic activity on the small molecular substrate *p*NPB. Multiple glass vials filled with 1 mL of 50 nM enzyme in glycine-NaOH buffer (50 mM, pH 9.2) were incubated at 40 °C in the absence or presence of 10 mg PET powders (LC-PET with 7.6% crystallinity or HC-PET with 30% crystallinity). The vials were taken after 1, 3, 6, 10, 15, 24, and 48 h incubation and cooled down to 4 °C within 10 min, then 20 μL of 0.2 mM pNPB in acetone was added to each vail for reaction. After 10-min hydrolysis at 4 °C, the absorbance at 405 nm of the solution was monitored on a Nanodrop Microvolume Spectrophotometer (Thermo Fisher Scientific Inc.). The concentration of the hydrolytic product *p*-nitrophenol was calculated using a molar extinction coefficient of 18,000 M^-1^ cm^-1^. Error bars were obtained from three measurements using parallel samples.

### AFM‐SMFS measurements

The AFM-SMFS measurement was performed on a Cypher VRS AFM (Oxford Instruments, UK) according to the protocol described previously^51^. Briefly, a silicon nitride AFM probe was cleaned with UV/Ozone treatment for 30 min and then silanized with a 3% (w/v) solution of 3-aminopropyltriethoxysilane (APTES) in ethanol for 1 h at room temperature. After drying at 110 °C for 30 min, the AFM tip was functionalized in sequence with 2.0 mg mL^-1^ NHS-PEG-MAL in dimethylformamide (DMF), 2.0 mg mL^-1^ HS-NTA in DMF, and 50 mM NiSO_4_ solution at room temperature. Each step of functionalization takes 1 hour. The enzyme grafting was achieved by immersing the functionalized AFM tip in 14 μM *Is*PETase or HotPETase for 1 h at room temperature. For measuring the adhesive force between the enzyme and a PET film, the cantilever’s spring constant was calibrated to be 0.06 N m^-1^. The enzyme-functionalized tip was moved toward a PET film (thicknesses of 0.25 mm) soaked in glycine-NaOH buffer (50 mM, pH 9.2) until they were in contact at a force of 0.5 nN for 0.5 s. Then the tip was retracted from the substrate at a 2.0 μm s^-1^ pulling speed.

## Data availability

The data that support the findings of this study are available from the corresponding author on reasonable request.

## Acknowledgments

This work was supported by the National Nature Science Foundation of China under grant number 32371325, and the seed funding of China Petrochemical Corporation (Sinopec Group) under grant number 223260.

## Author Contributions

Y.C., S.H., and Y.Z. supervised the project. Y.C. conceived the idea, built the model, and analyzed the results. S.C. performed all the MD simulations and analysis of the results. E.A. carried out the experiments. W.Q. assisted in data analysis. Y.C., S.C., S.H., and Y.Z. co-wrote the manuscript. All authors discussed the results and assisted with the manuscript preparation.

