## supplementary material for "A Mechanistic Understanding of the Activity-Stability Trade-off in PETase"

**Supplementary Note 1**

We speculated that the flexible loop is necessary for a PETase with high activity, as it enhances the enzyme's ability to capture PET fragments. *Is*PETase exhibits a highly flexible loop at Ile208 and wobbling at Trp185, which adopt a confined conformation in the PET-bound state (Fig. 2d-e, Supplementary Fig. 17-21). This dual behavior—flexibility in the unbound state and a tightly confined conformation in the PET-bound state—is the origin of *Is*PETase's high PET depolymerization activity at moderate temperatures. This conclusion is supported by a previous study by Rey-Ting Guo[1], where H218S and F222I mutations were introduced into LCC (LCC-DM) to induce Trp185 wobbling. The PET depolymerization activity of LCC at low temperatures increased, though it showed lower activity at high temperatures. Our simulations confirmed that the wobbling of the Trp residue was successfully introduced into LCC-DM, and LCC-DM exhibited higher loop flexibility at 300 K (Supplementary Fig. 22-23). However, LCC-DM was unstable at high temperatures (353.15 K), and the conformation of the catalytic triad could be disrupted. In contrast, wild-type LCC remained stable at high temperatures. These results further confirm the activity-stability trade-off in *Is*PETase.

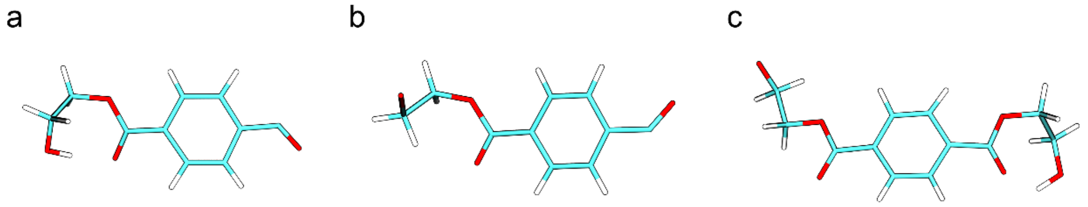


**Supplementary Figure 1.** The (a) head, (b) repeat unit, and (c) tail of the polyethylene terephthalate (PET) used in our simulations.

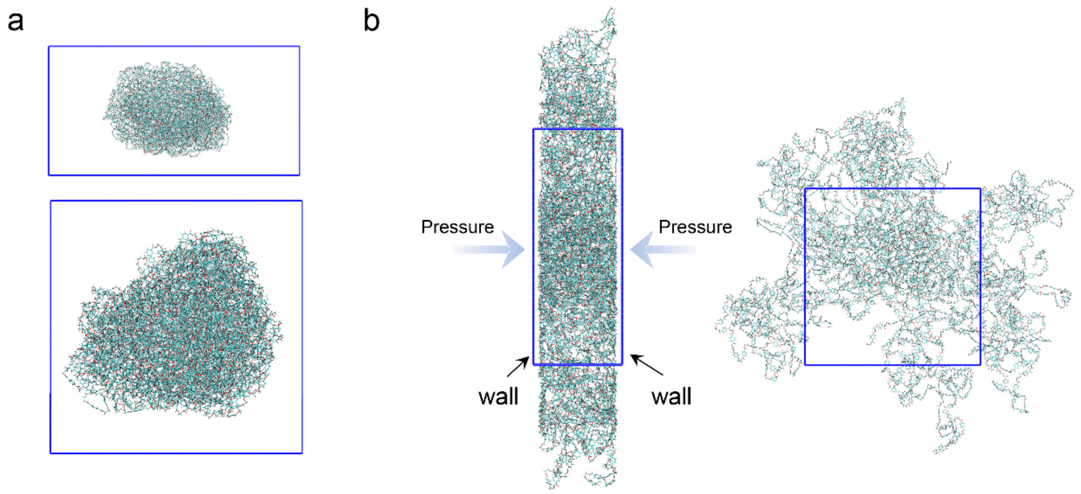


**Supplementary Figure 2.** The construction of PET membrane model for interface simulation. (a) The initial state of 12 PET chains. (b) The x and y directions of the simulation box have periodic boundary conditions, while z=0 and z=box positions are occupied by two walls composed of CA atoms (atom type in Amber force field). Pressure is applied in the z direction, causing PET to compress into a membrane.

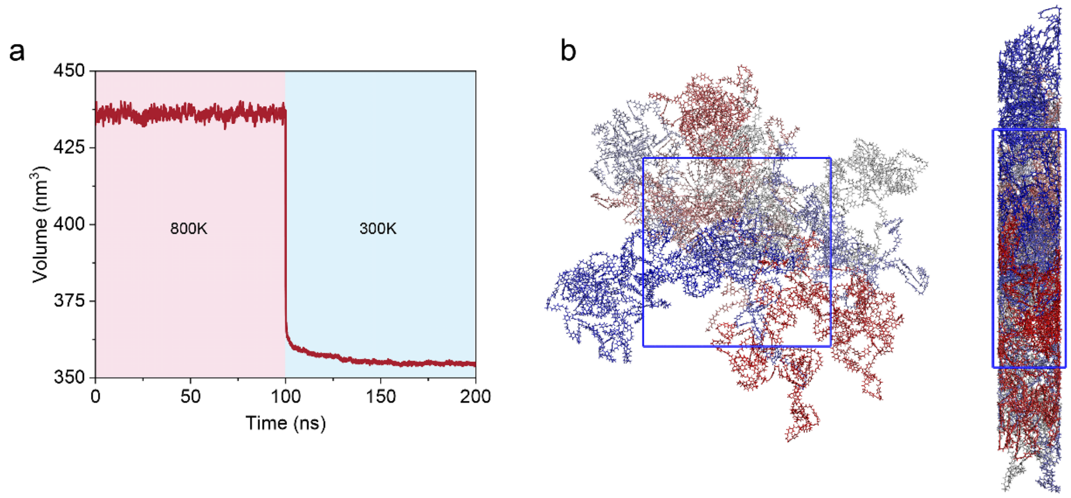


**Supplementary Figure 3.** PET membrane simulation. To achieve homogeneity of different PET chains in the membrane, we first simulated for 100 ns at 800 K, followed by another simulation for 100 ns at 300 K. (a) The variation in box volume during MD simulations. (b) Visualization of the polymer conformation of PET within the membrane, with different chains depicted in different colors.

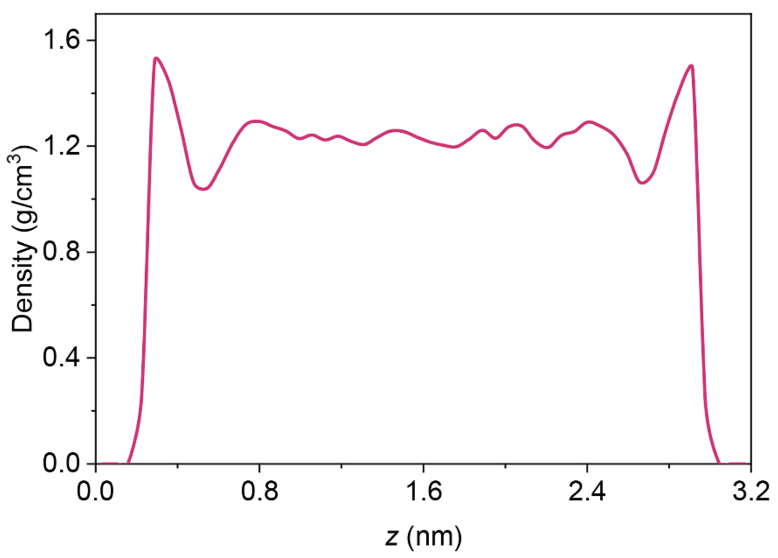


**Supplementary Figure 4.** The density profile of the PET membrane along the z-axes of the simulation box.

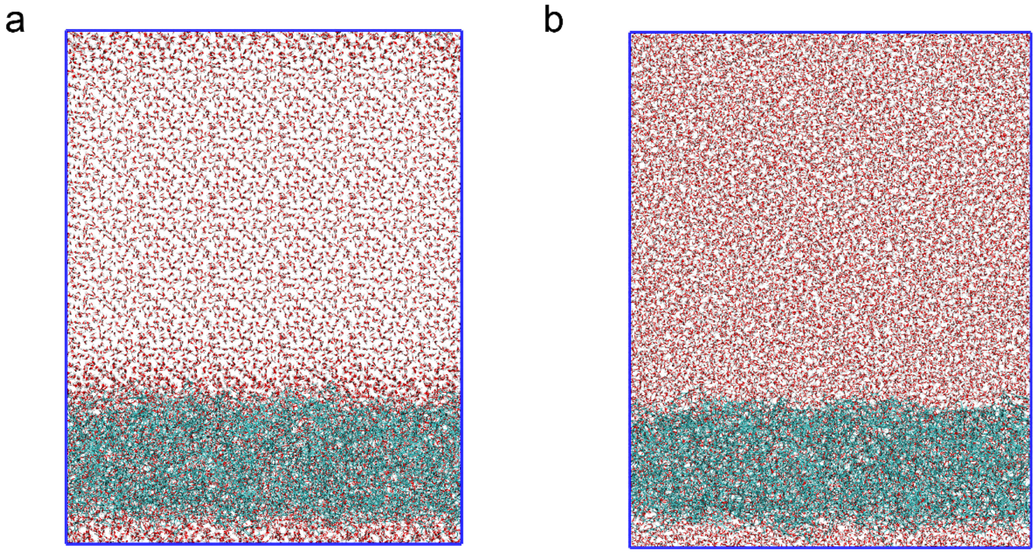


**Supplementary Figure 5.** The solid-liquid interface model of PET in simulations: (a) depicts the initial state, while (b) illustrates the equilibrium state of the model.

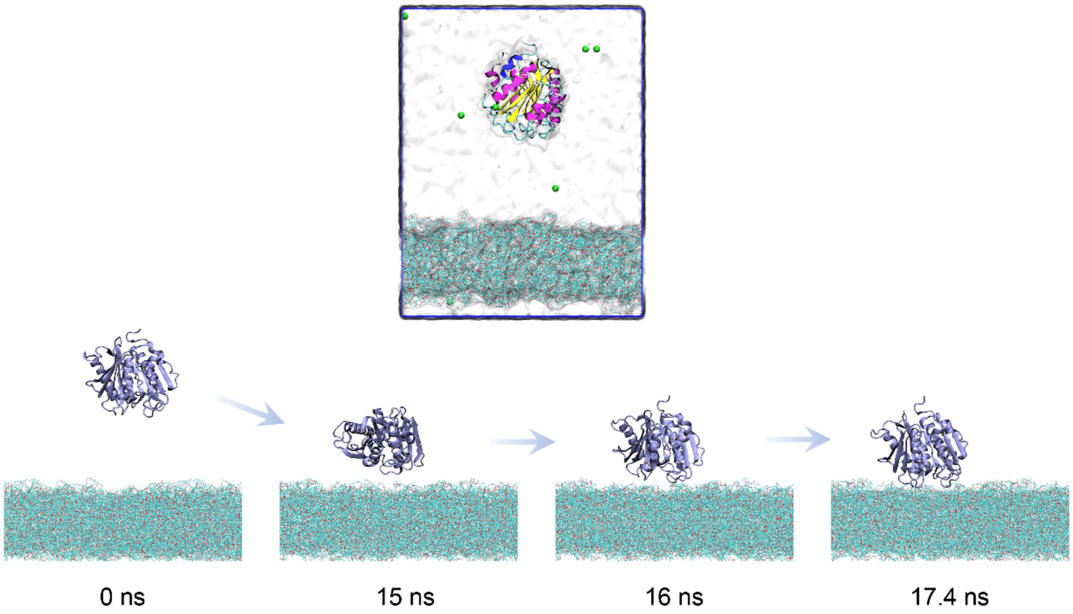


**Supplementary Figure 6.** The supervised MD simulation aims to sample the binding conformations of the PETase-PET complex. The top figure depicts the initial states of the simulation, with the enzyme positioned sufficiently away from the PET surface. The bottom figure illustrates the binding dynamics of PETase on the PET surface.

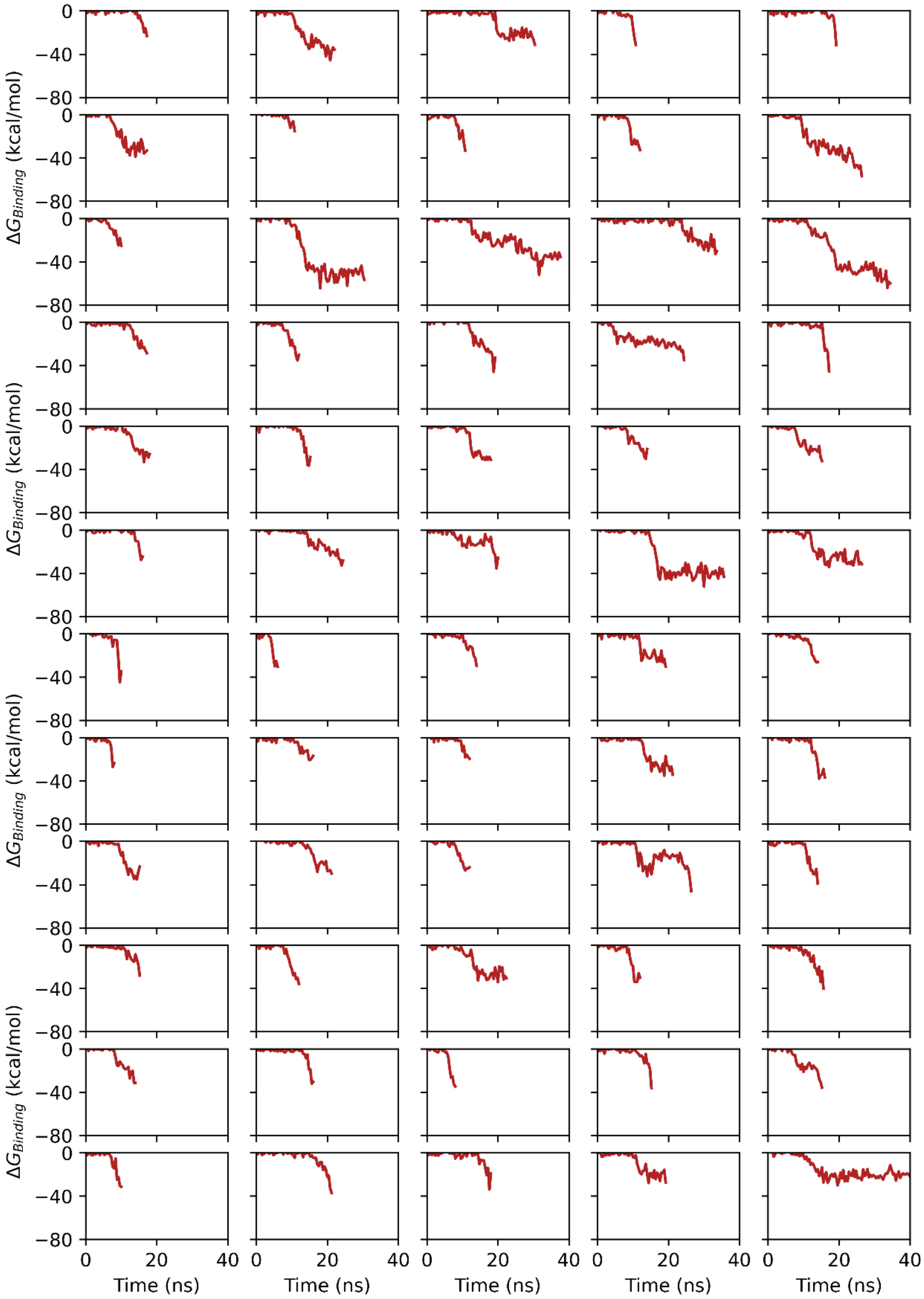


**Supplementary Figure 7. (a)** The binding free energy between the enzyme and the PET surface was calculated using MM/PBSA during supervised MD simulations. Each subfigure depicts the evolution of binding energy during one sampling. Results from samplings 1-60 are displayed.

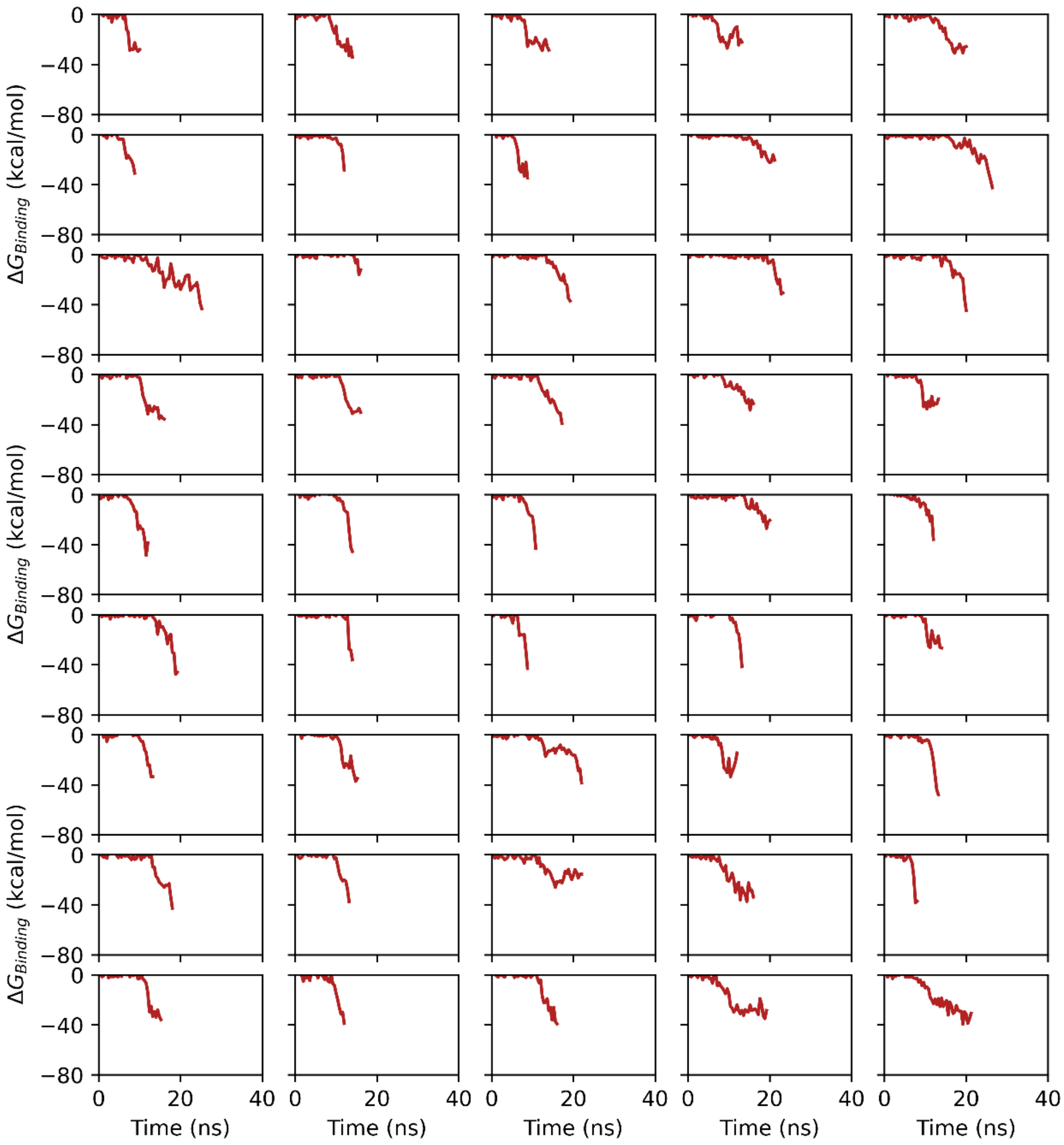


**Supplementary Figure 7. (b)** Results from samplings 61-105 are displayed.

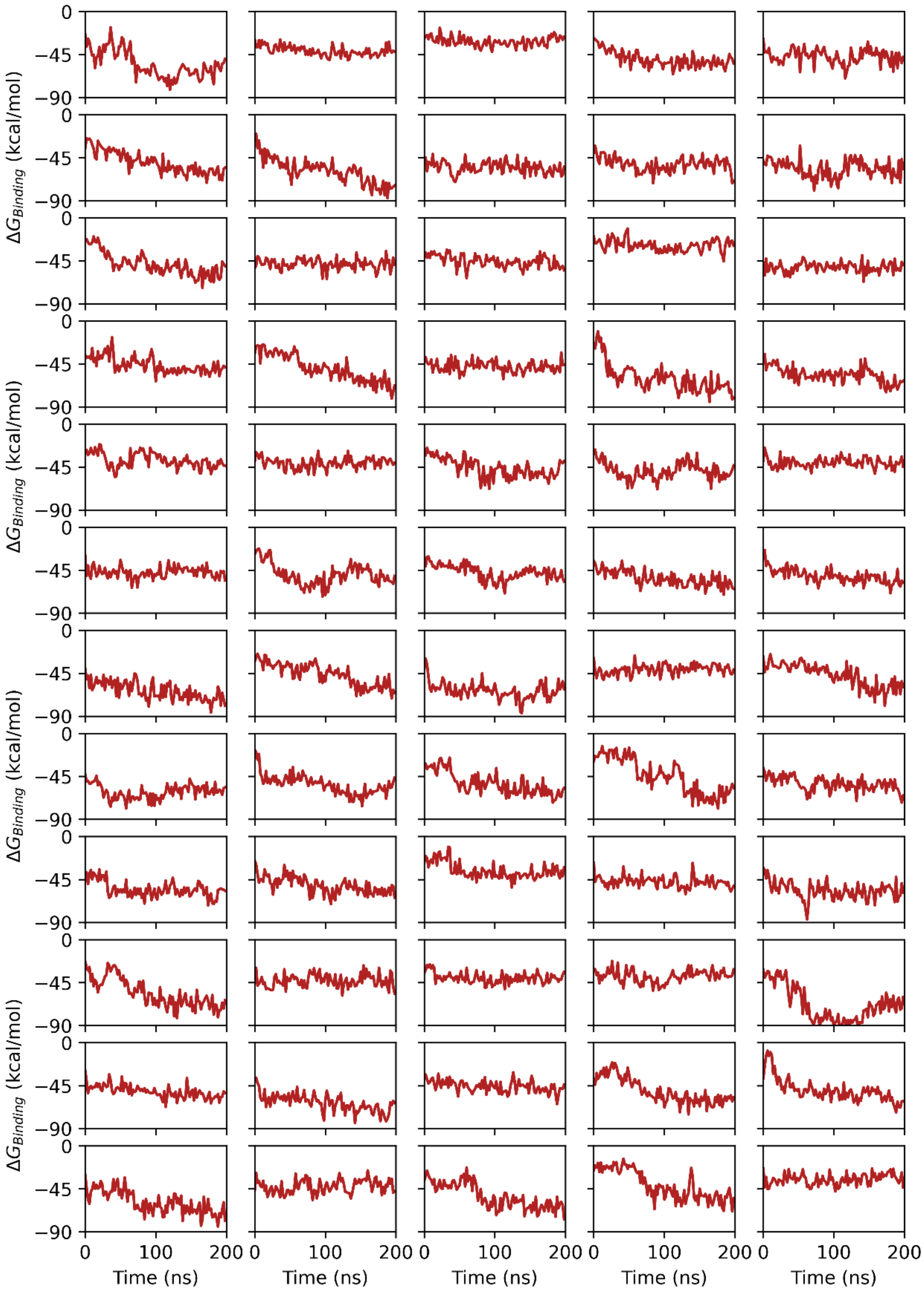


**Supplementary Figure 8. (a)** The binding free energy (MM/PBSA) during extended 200 ns MD simulations. Results from samplings 1-60 are displayed.

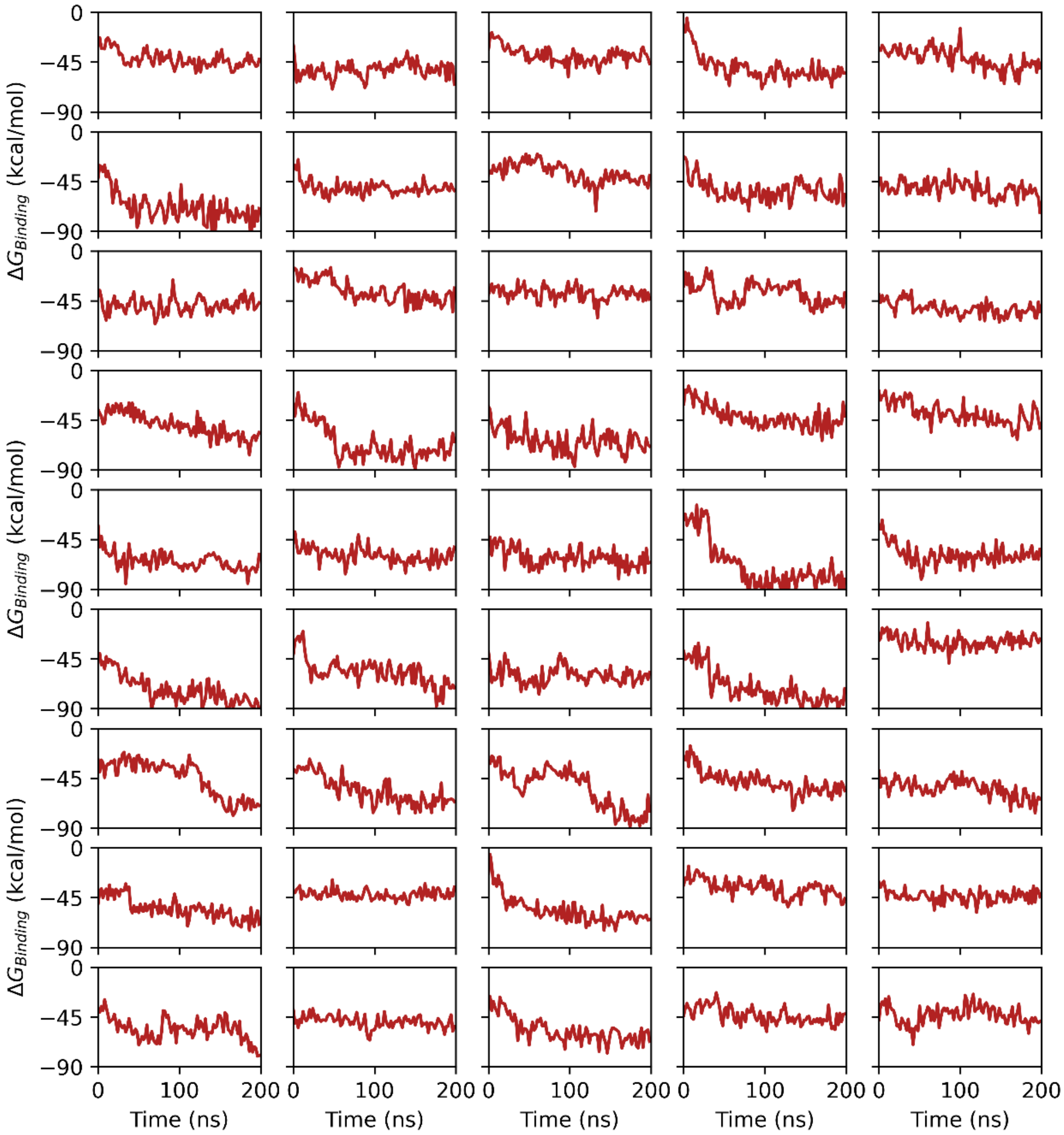


**Supplementary Figure 8. (b)** Results from samplings 61-105 are displayed.

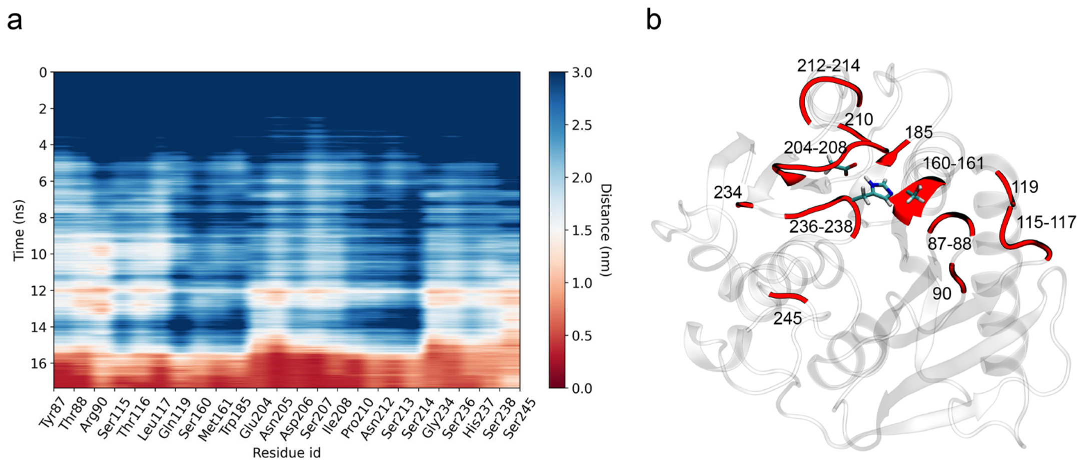


**Supplementary Figure 9.** Distances between some key residues of *Is*PETase and PET during supervised MD. (a) The evolution of distances throughout the simulation. (b) The location of the residues listed in (a) on the enzyme surface.

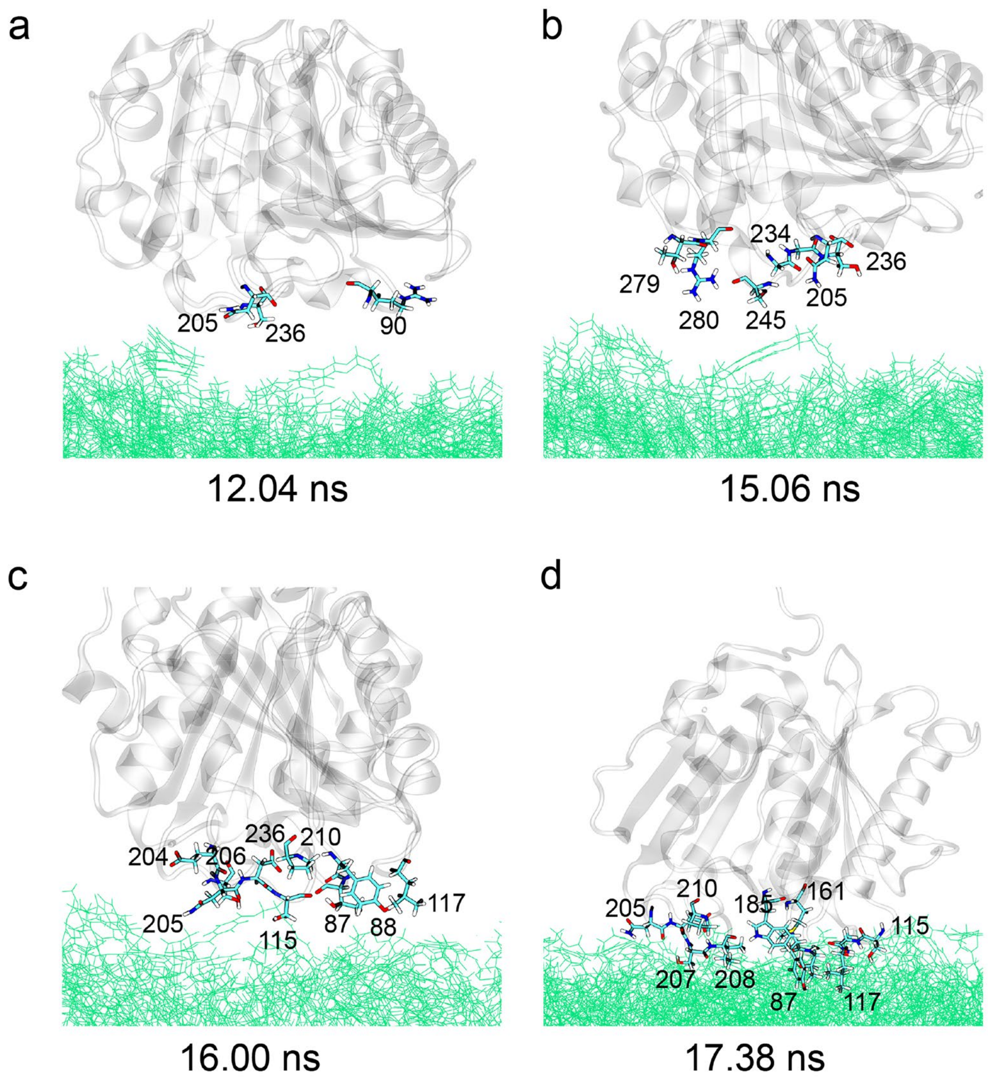


**Supplementary Figure 10.** Residues in proximity to the PET surface during supervised MD. They include Tyr87, Thr88, Arg90, Ser115, Leu117, Met161, Trp185, Glu204, Asn205, Asp206, Ser207, Ile208, Pro210, Gly234, Ser236, Ser245, Thr279, and Arg280. The distance cut-offs in a-d are 9, 7.5, 5, and 3.5 Å, respectively.

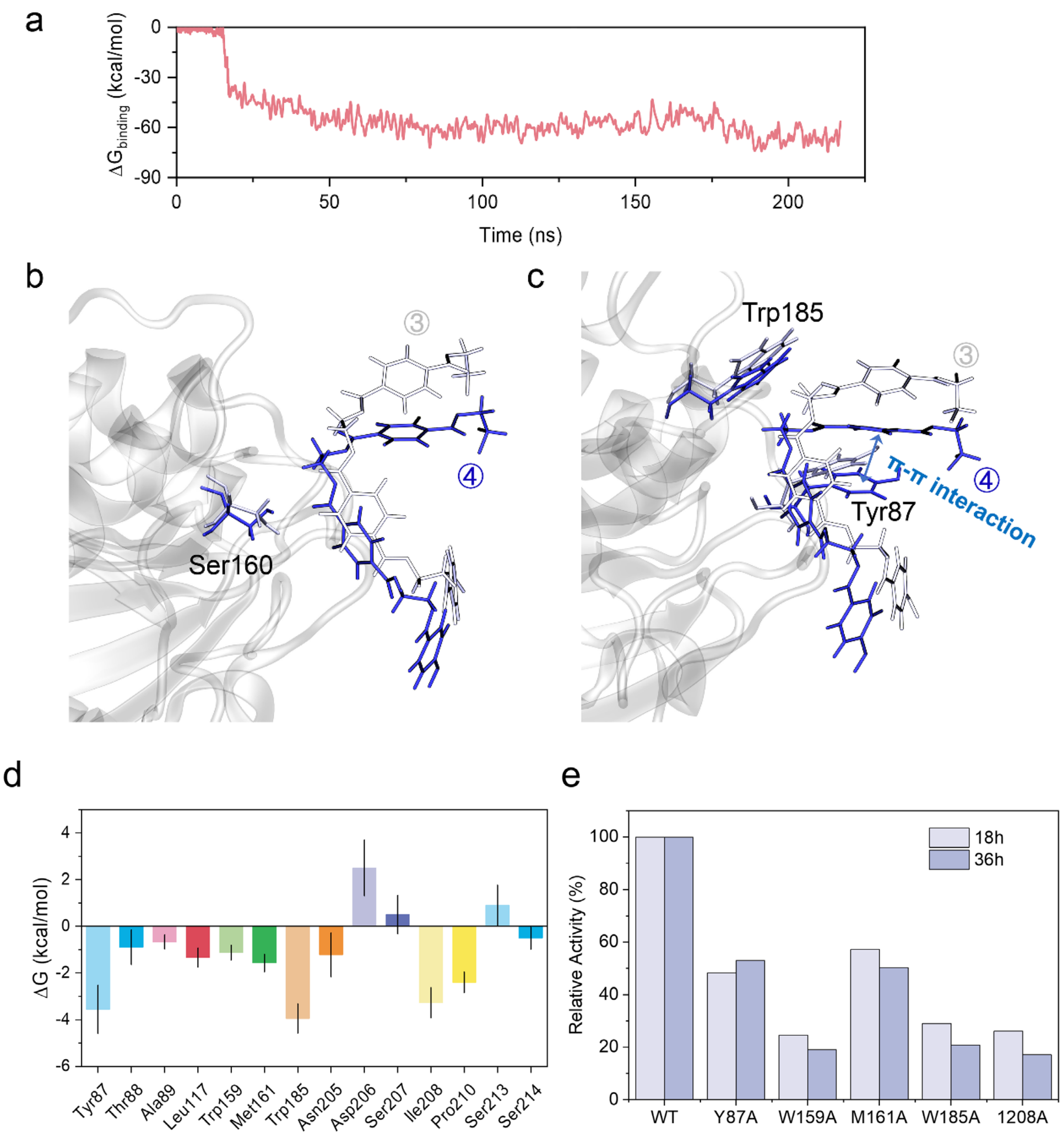


**Supplementary Figure 11.** (a) Binding free energy of *Is*PETase during supervised and extended (200 ns) molecular dynamics (MD) simulations. (b) The captured PET fragment exhibited close proximity after refinement of the binding conformation (③→④). The ③ and ④ represent the conformational refinement and pre-cleavage states in Fig. 1d. (c) Following the conformational refinement (③→④), the PET fragment formed a Π-Π interaction with Tyr87. (d) The top 14 residues of *Is*PETase contributing the most to the binding free energy with PET fragments. (e) The PETase activity of wild-type and mutant *Is*PETase was assessed using PET film as a substrate. Data extracted from Joo et al[2].

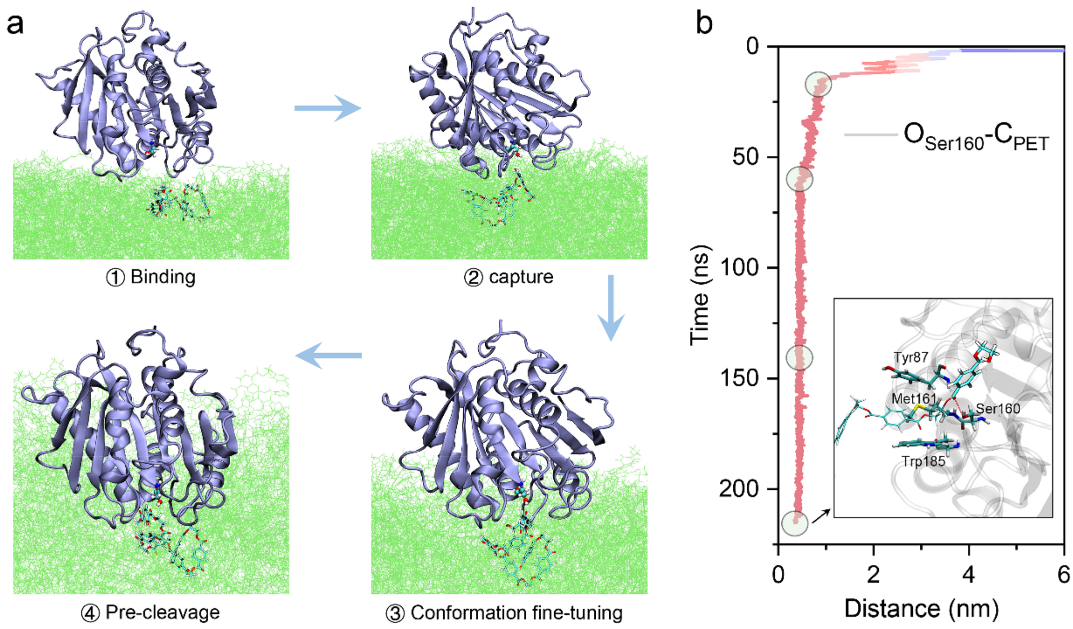


**Supplementary Figure 12. Ester bond recognition by *Is*PETase in an exolytic mode.** (a) Detailed mechanism of ester bond recognition by *Is*PETase. (b) Distance between the oxygen atom of the side chain hydroxyl of Ser160 and the carbon atom of the recognized PET ester bond.

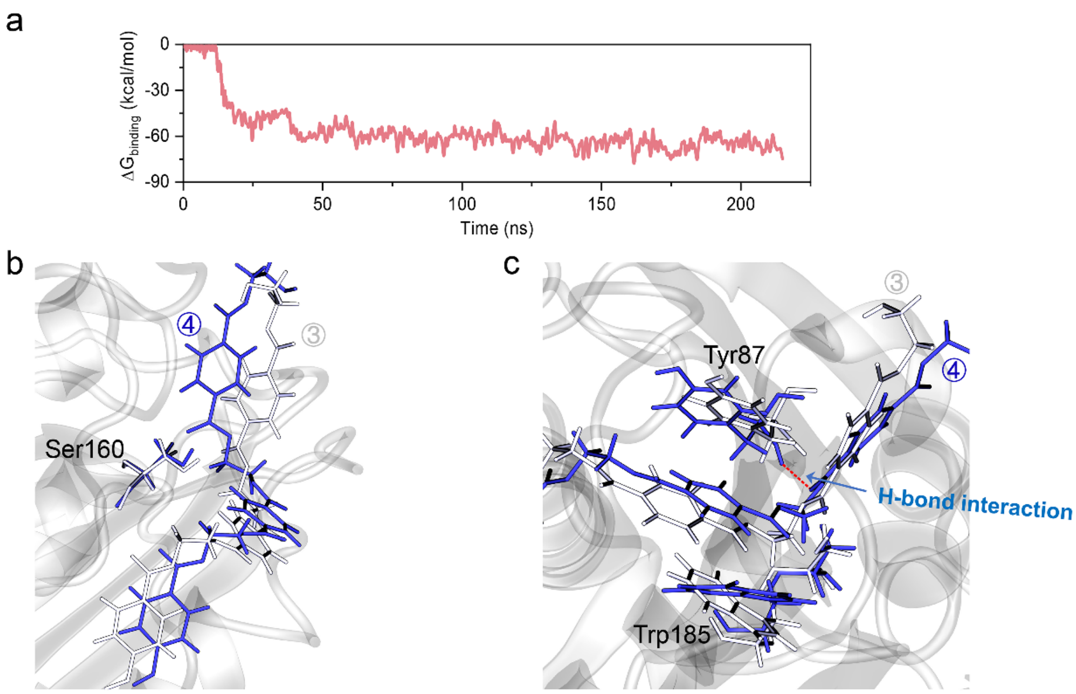


**Supplementary Figure 13.** (a) Binding free energy of *Is*PETase (exolytic mode) during supervised and extended (200 ns) molecular dynamics (MD) simulations. (b) The captured PET fragment exhibited close proximity after refinement of the binding conformation (③→④). The ③ and ④ represent the conformational refinement and pre-cleavage states. (c) Following the conformational refinement (③→④), the PET fragment formed a H-bond interaction with Tyr87.

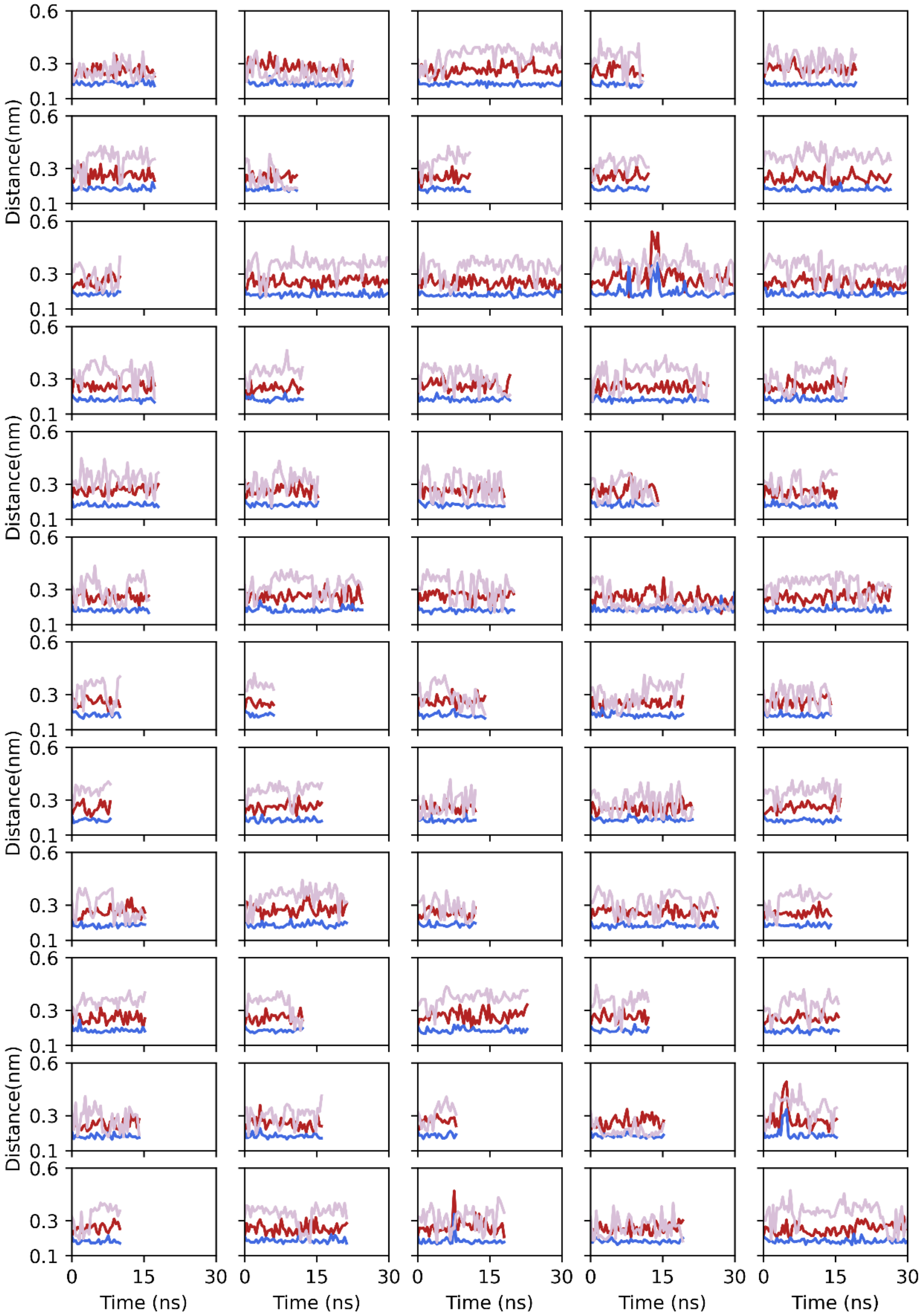


**Supplementary Figure 14.** (a) Three pairs of distances within the enzyme active site during supervised MD simulations of *Is*PETase. Each subfigure depicts the evolution of distances during one sampling. Results from samplings 1-60 are displayed.

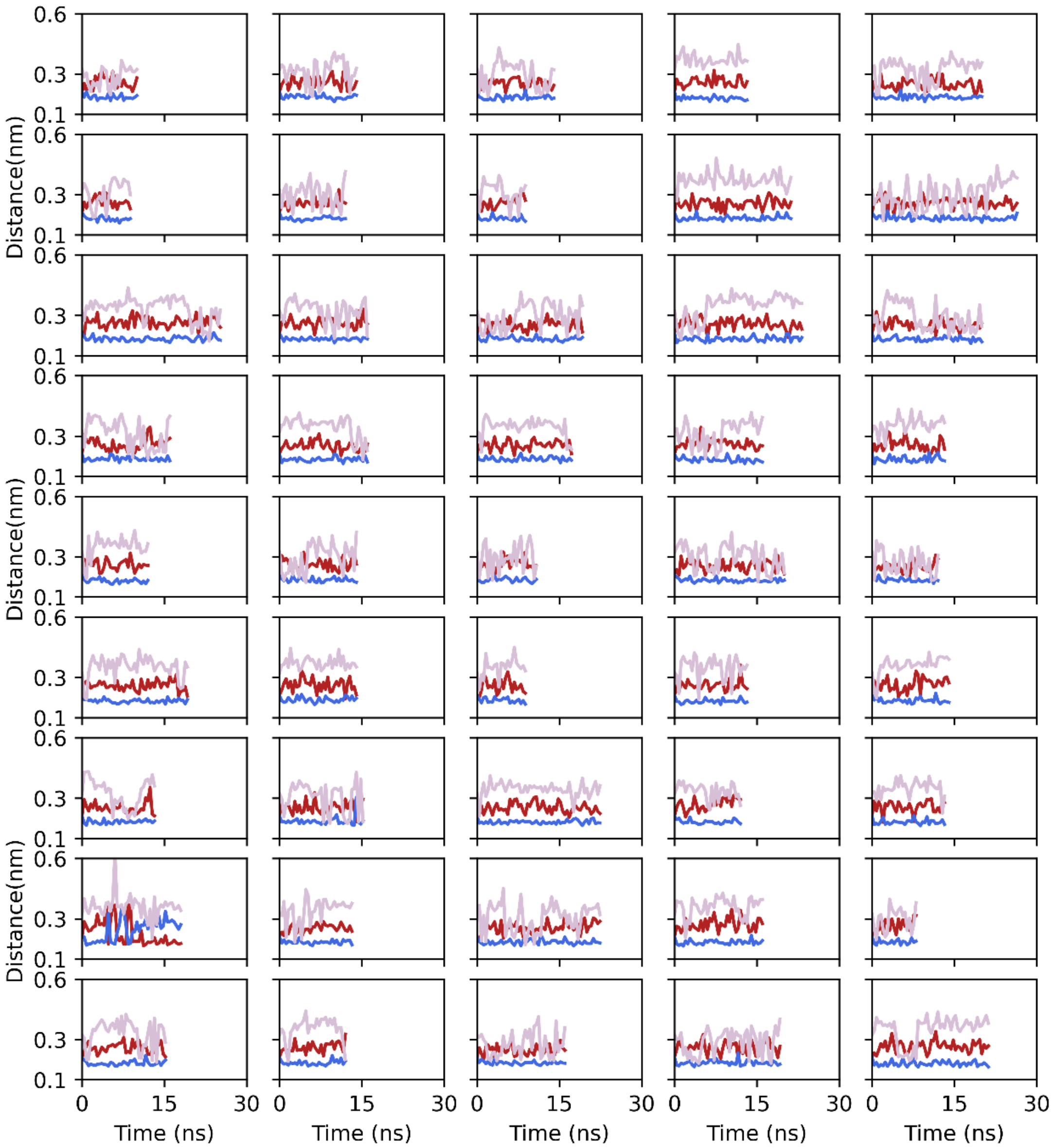


**Supplementary Figure 14.** (b) Results from samplings 61-105 are displayed.

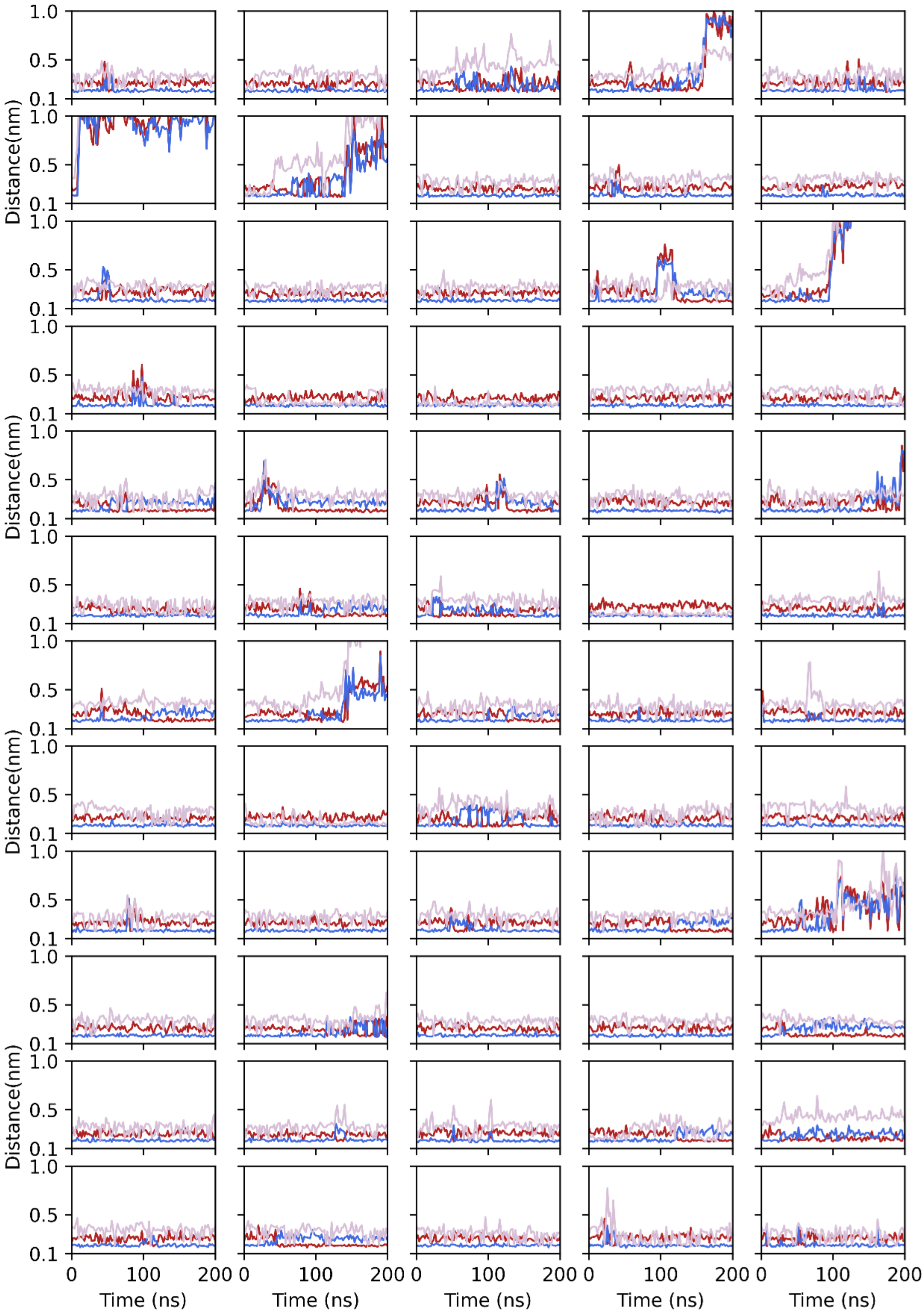


**Supplementary Figure 15.** (a) Three pairs of distances within the enzyme active site during 200 ns extended MD simulations of *Is*PETase. Results from samplings 1-60 are displayed.

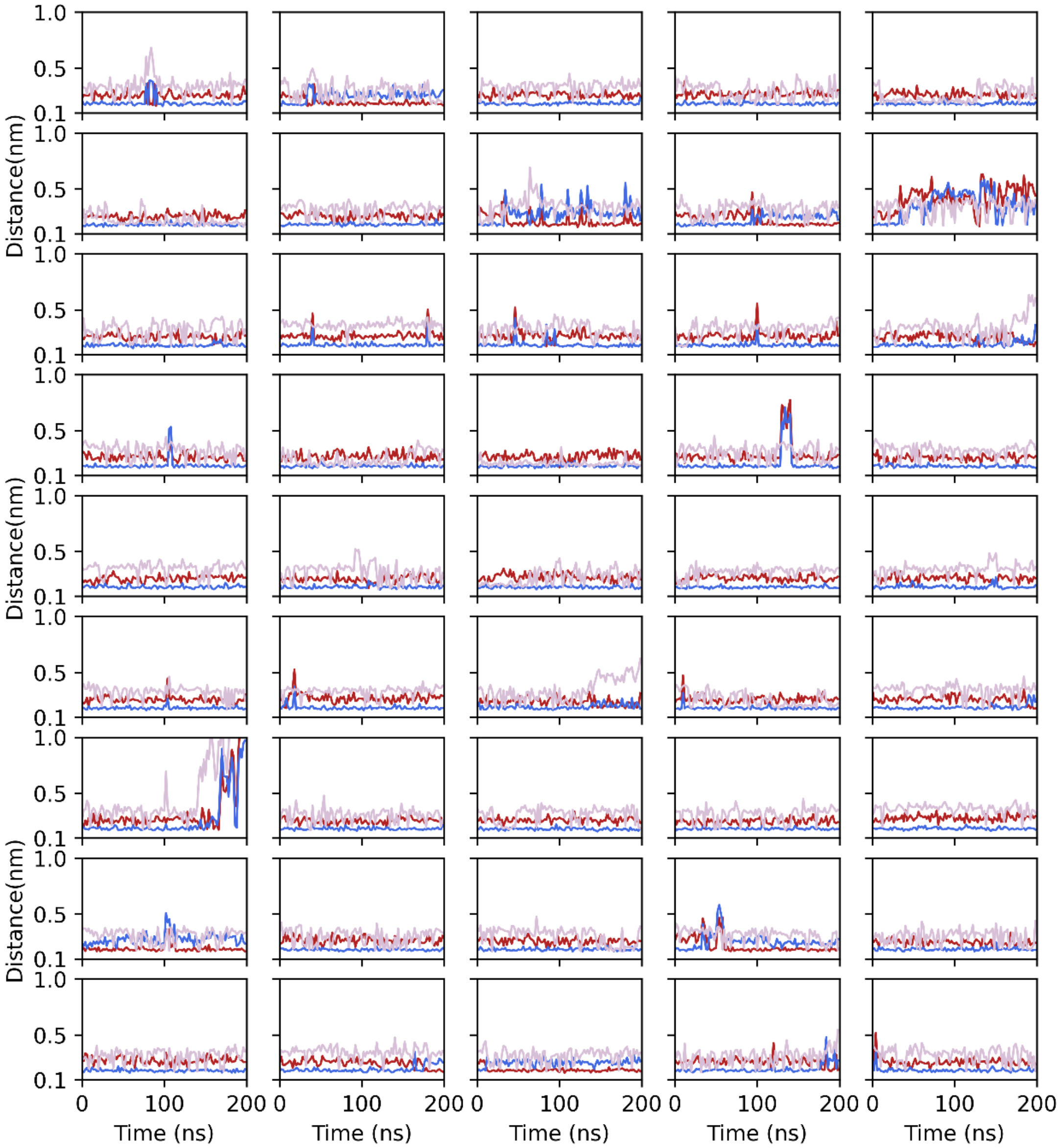


**Supplementary Figure 15.** (b) Results from samplings 61-105 are displayed.

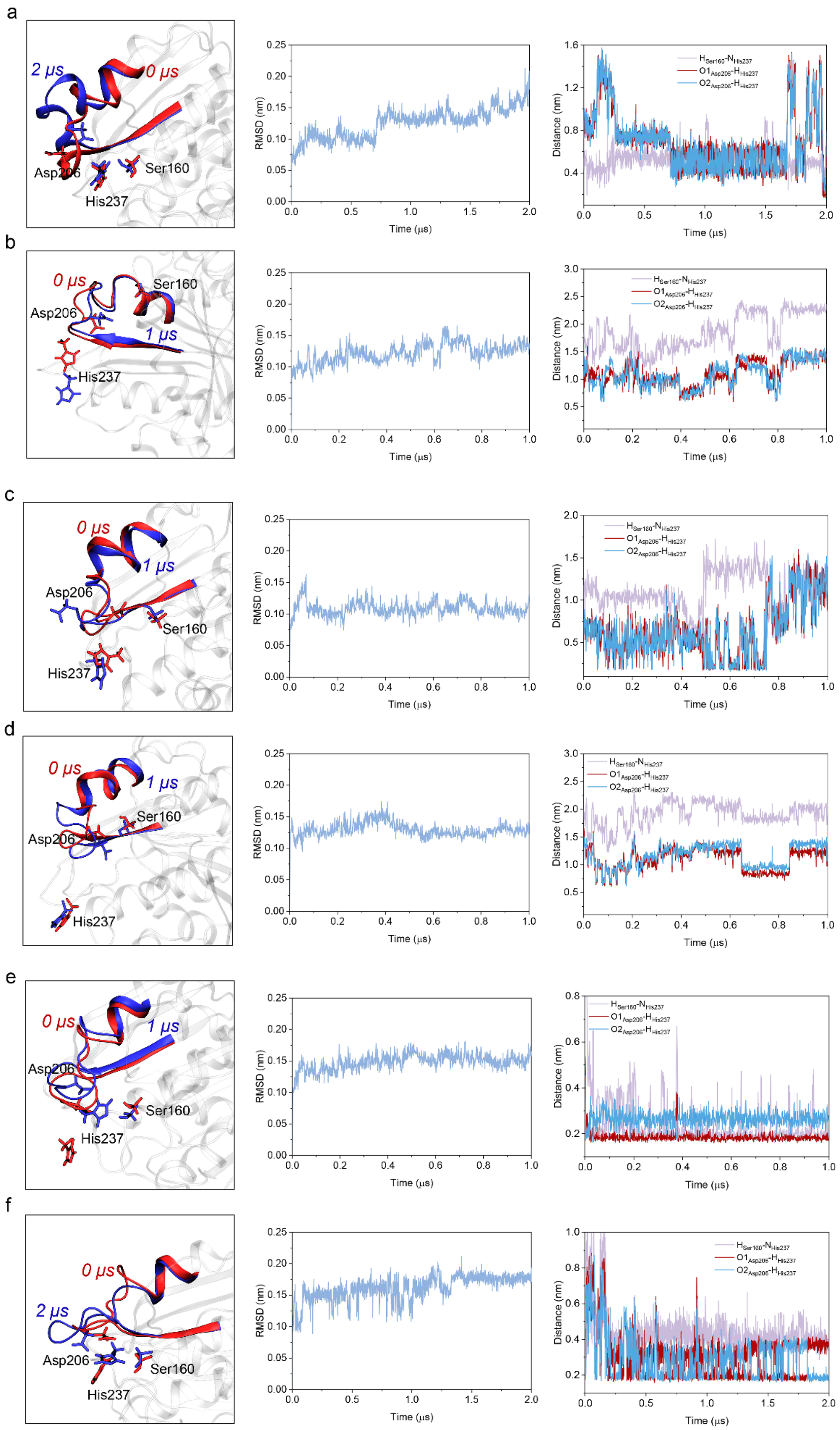


**Supplementary Figure 16.** Long time MD simulations (1 μs or 2 μs) for disrupted states of *Is*PETase. The initial conformations of enzyme in subfigures a-f were taken from samplings 4, 6, 7,

15, 32, and 45 in Supplementary Figure 15a. Throughout the MD simulations, the enzyme remained unbound in water. In subfigures a-d, the conformation of the enzyme's active site remained deactivated, indicating that the enzyme may have been permanently deactivated. In subfigures e-f, the enzyme's active state was restored.

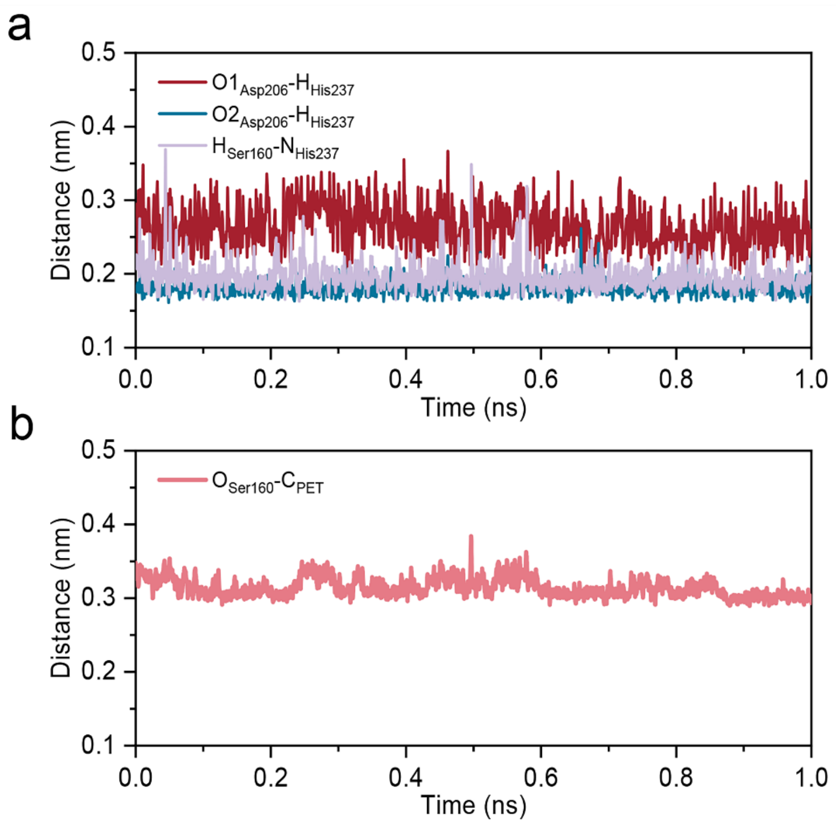


**Supplementary Figure 17. 1 μs MD simulation of *Is*PETase-PET complex (endolytic mode)**. (a) Three pairs of distances within the active site of *Is*PETase. (2) Distance between the oxygen atom of the side chain hydroxyl of Ser160 and the carbon atom of the recognized PET ester bond.

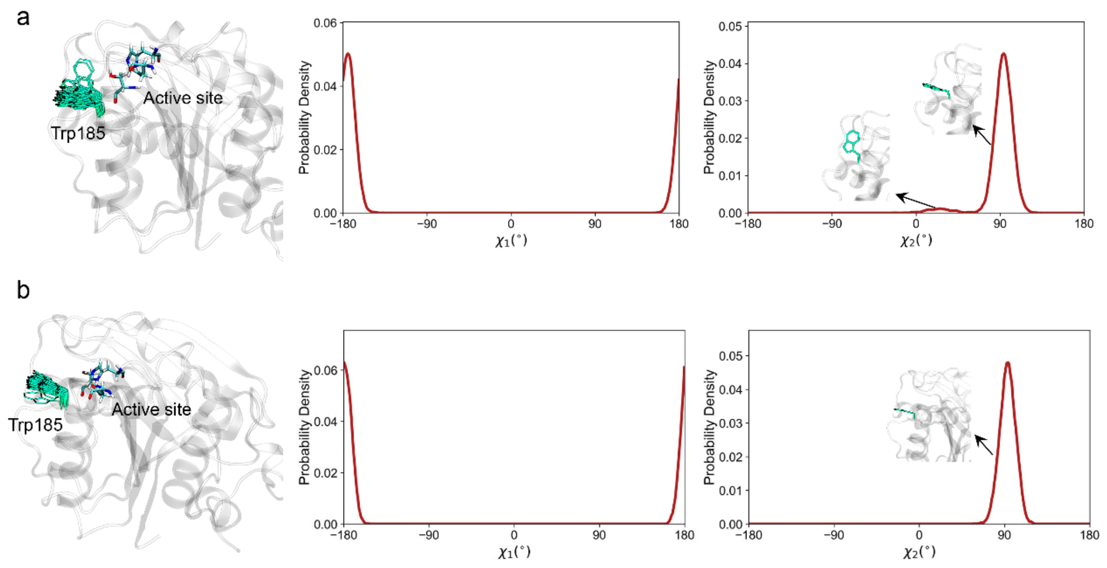


**Supplementary Figure 18.** Side chain torsion angle distribution of Trp185 in *Is*PETase and PET-bound *Is*PETase. Conformation of Trp185 in (a) *Is*PETase, (b) PET-bound *Is*PETase, and probability distribution of $\chi_1$ and $\chi_2$.

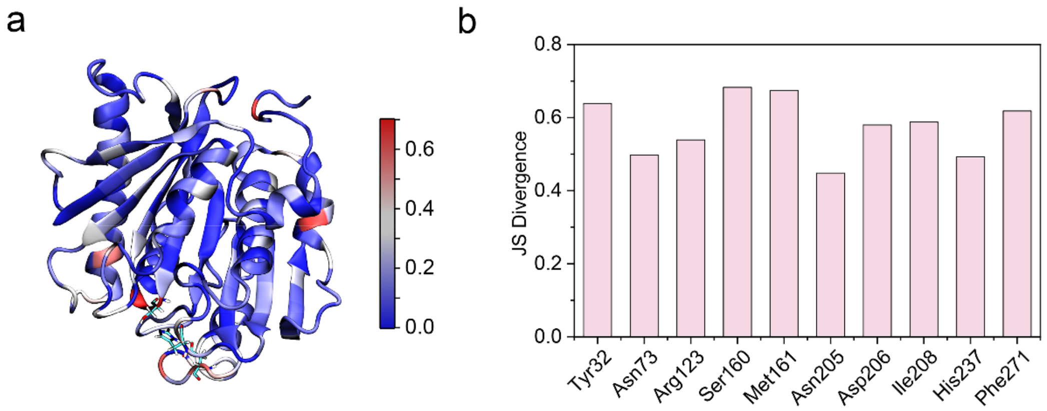


**Supplementary Figure 19.** (a) Visualization of JS divergence of the probability distribution of side-chain torsion angles between PET-bound and free *Is*PETase. (2) The top 10 residues with the highest JS divergence values.

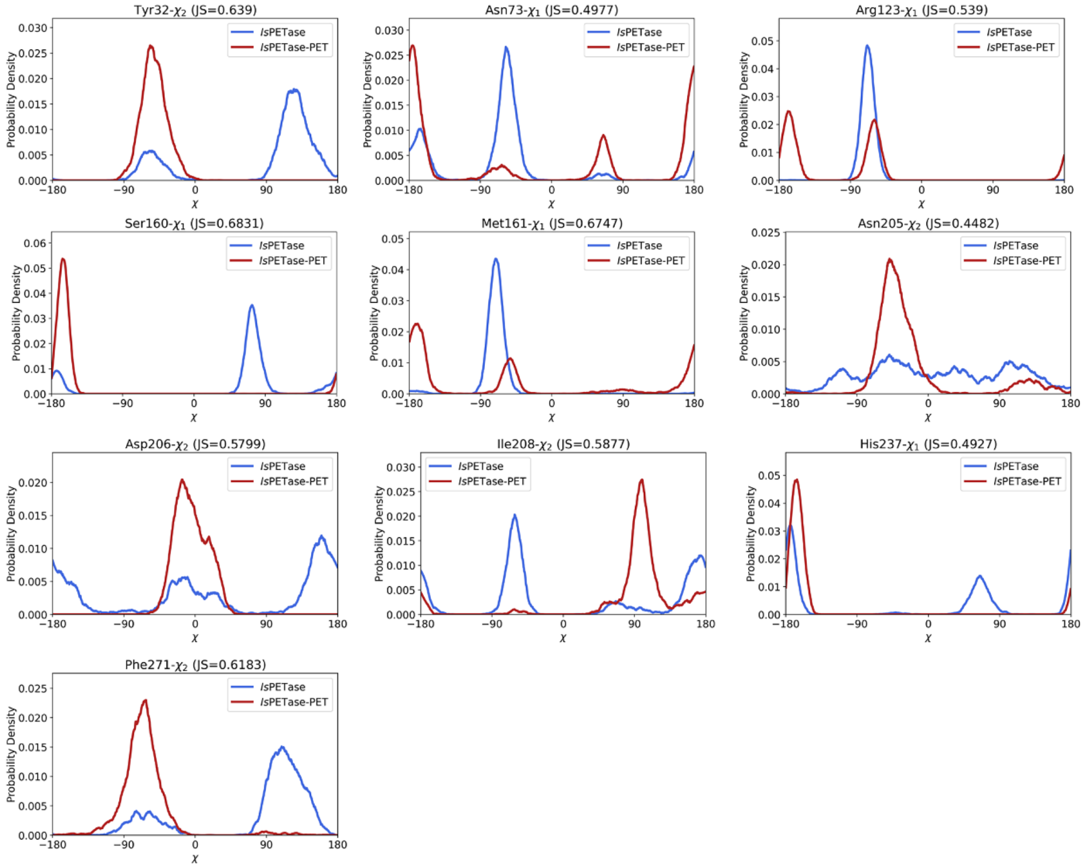


**Supplementary Figure 20.** Probability distribution of side-chain torsion angles of top 10 residues in Supplementary Figure 18.

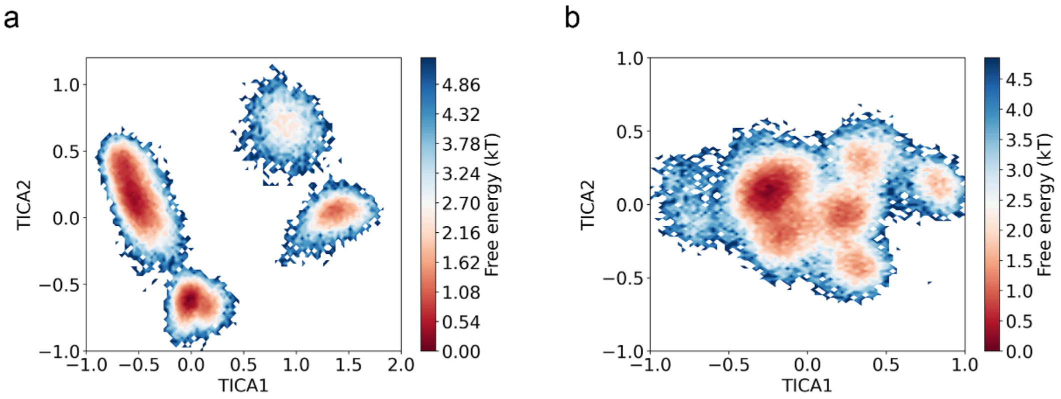


**Supplementary Figure 21.** The free energy landscape of (a) unbound and (b) PET bound *Is*PETase was constructed using the TICA method for dimensionality reduction. The features utilized for this analysis included the side-chain torsion angles of ten residues in Supplementary Figure 18.

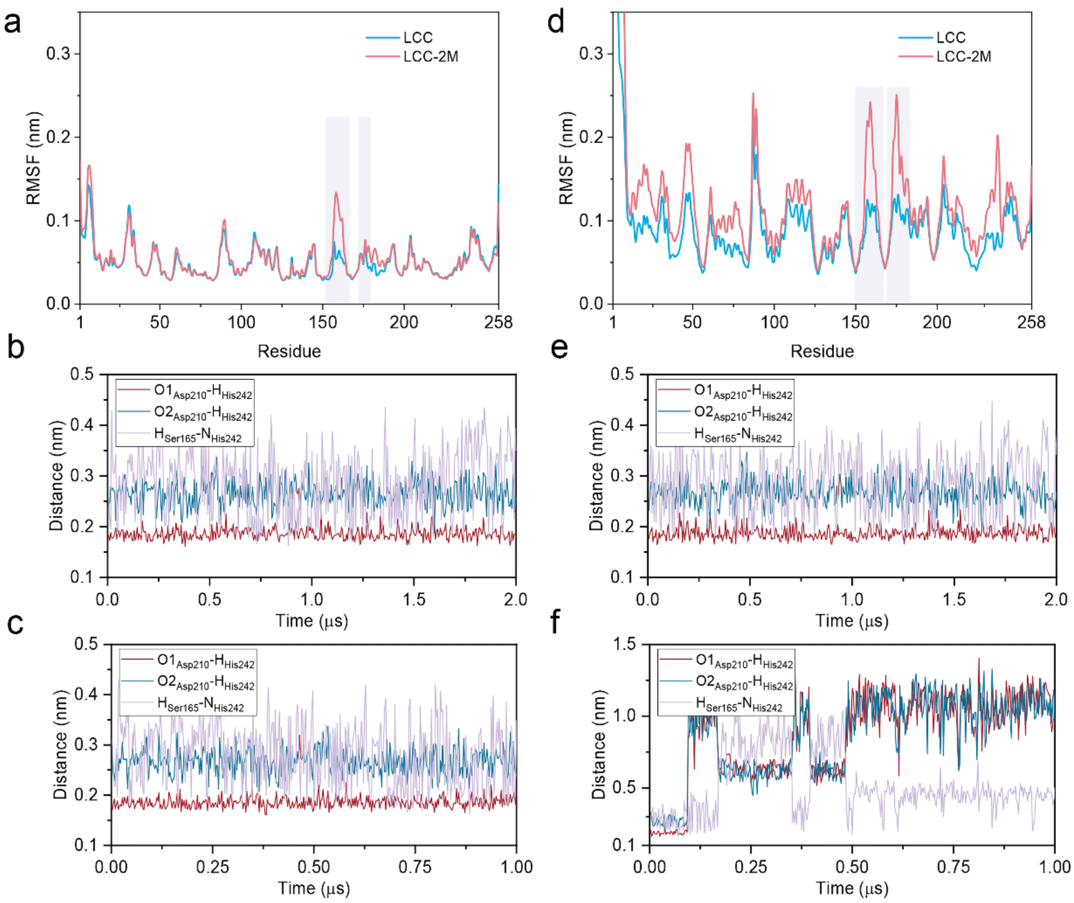


**Supplementary Figure 22.** The RMSF of LCC and LCC-DM (mutant H218S & F222I) at (a) 300 K and (d) 353.15 K. The distance pairs of the active site of LCC at (b) 300 K and (c) 353.15 K. The distance pairs of the active site of LCC-DM at (e) 300 K and (f) 353.15 K.

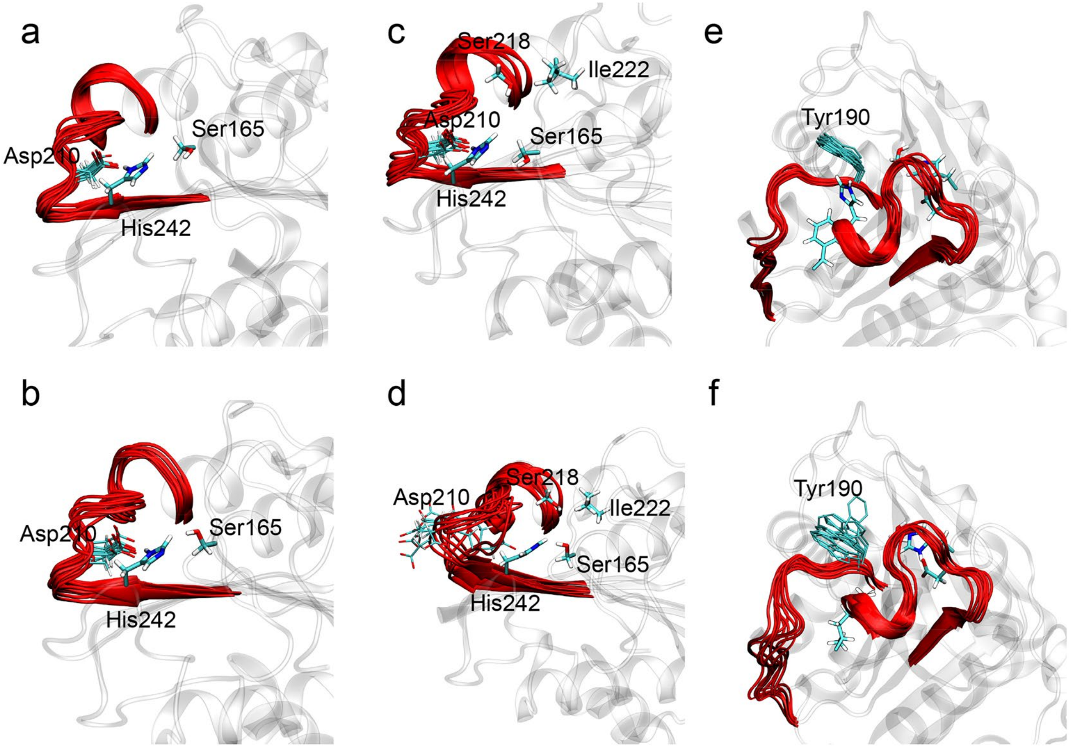


**Supplementary Figure 23.** Superposition of multi-frame structures: (a) LCC at 300 K, (b) LCC-DM (H218S and F222I) at 300 K, (c) LCC at 353.15 K, (d) LCC-DM at 353.15 K. The catalytic loop and catalytic triad were shown. (e) Conformation of W190 (W185 in *Is*PETase) in LCC at 300 K. (f) Wobbling of W190 in LCC-DM at 300 K.

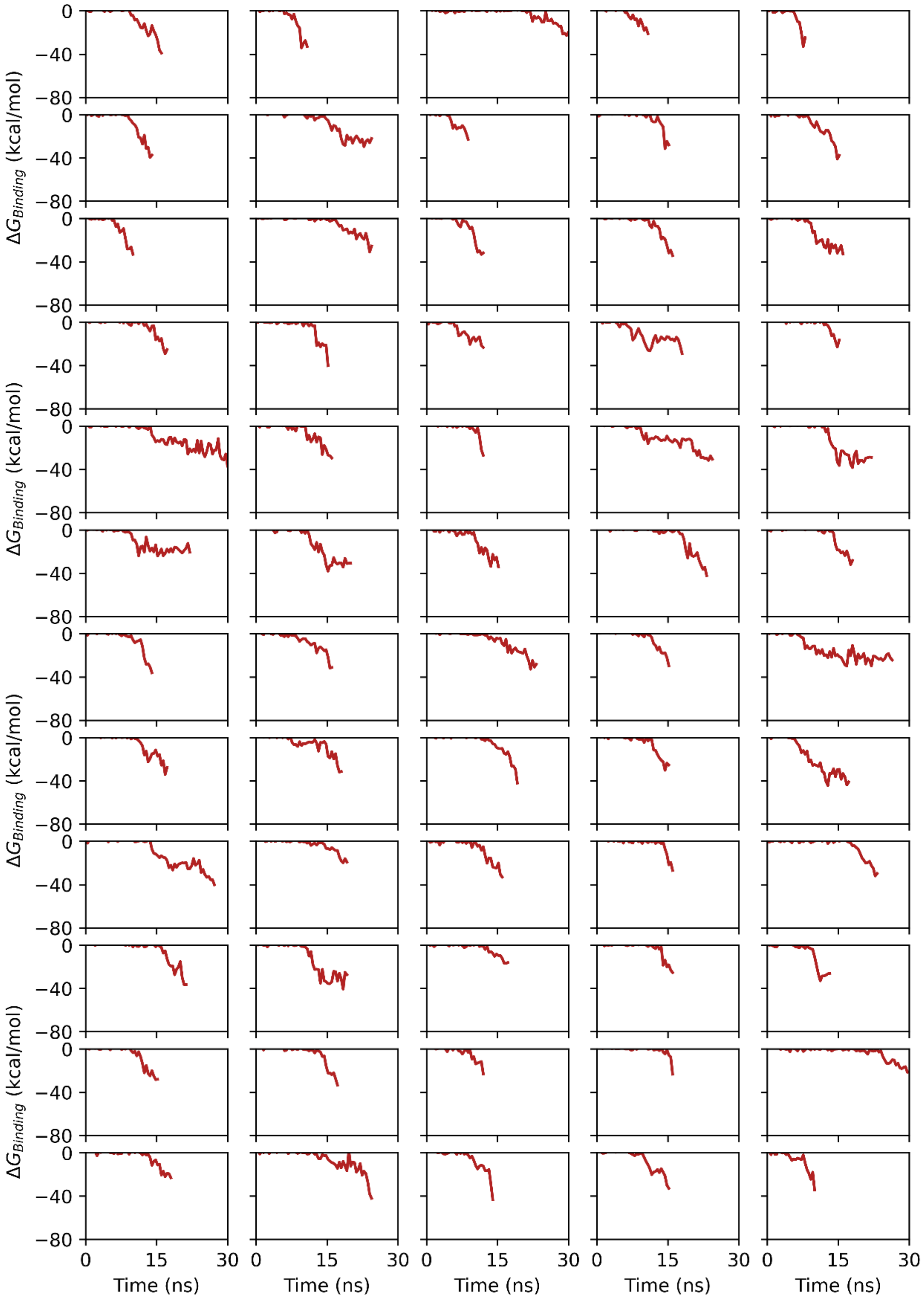


**Supplementary Figure 24. (a)** The binding free energy between the HotPETase and the PET surface was calculated using MM/PBSA during supervised MD simulations. Each subfigure depicts the evolution of binding energy during one sampling. Results from samplings 1-60 are displayed.

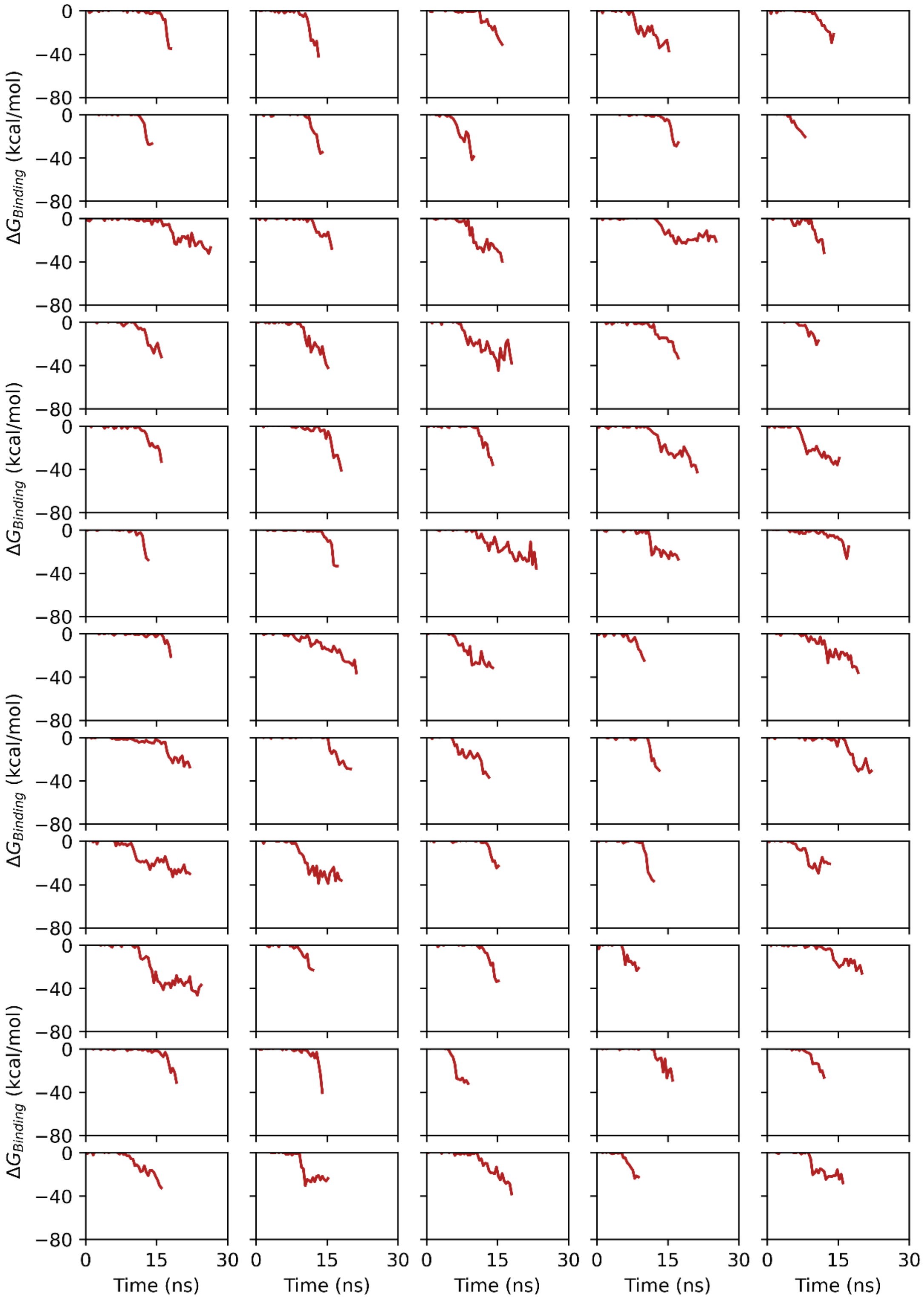


**Supplementary Figure 24. (b)** Results from samplings 61-120 are displayed.

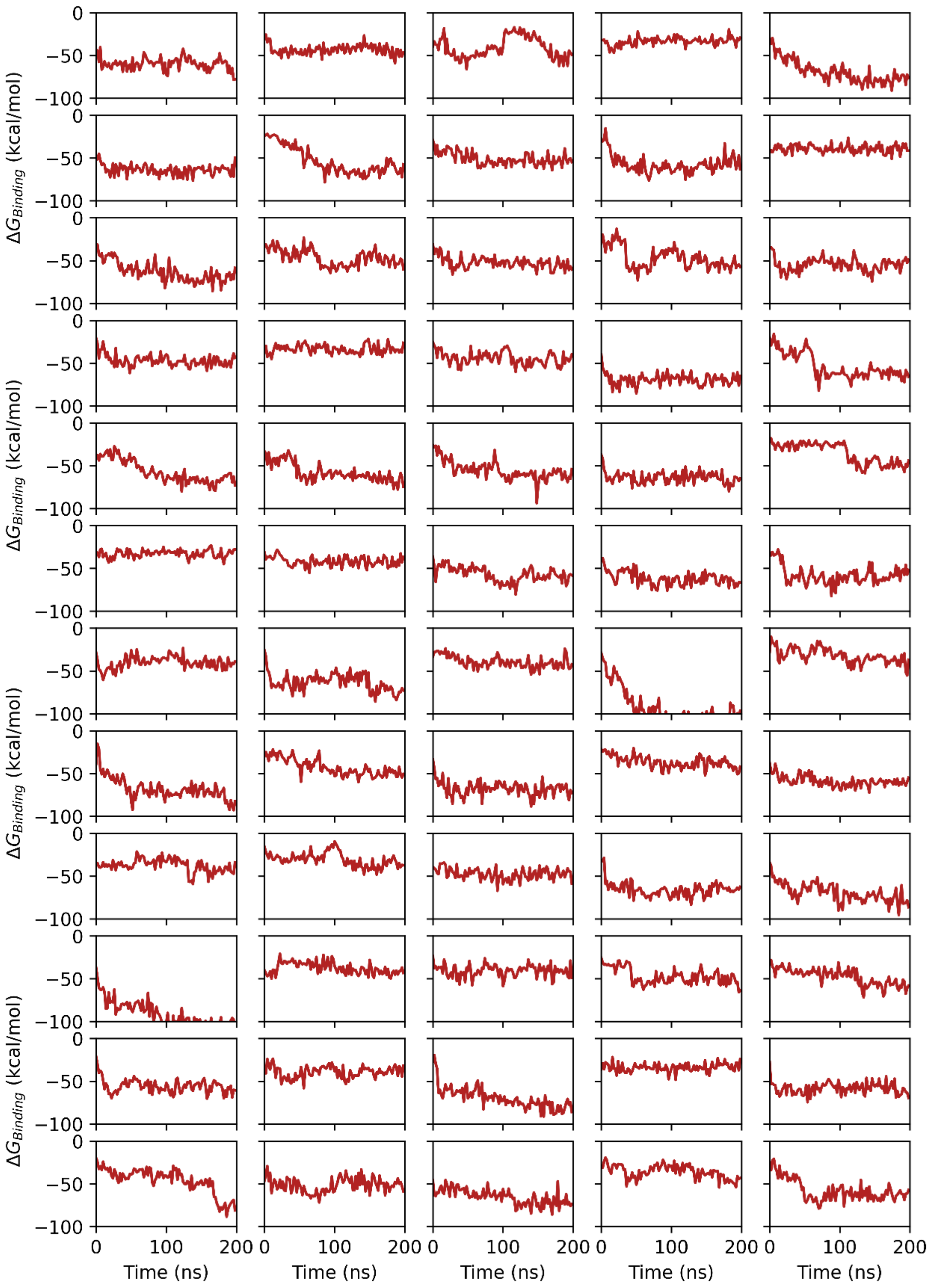


**Supplementary Figure 25. (a)** The binding free energy (MM/PBSA) during extended 200 ns MD simulations. Results from samplings 1-60 are displayed.

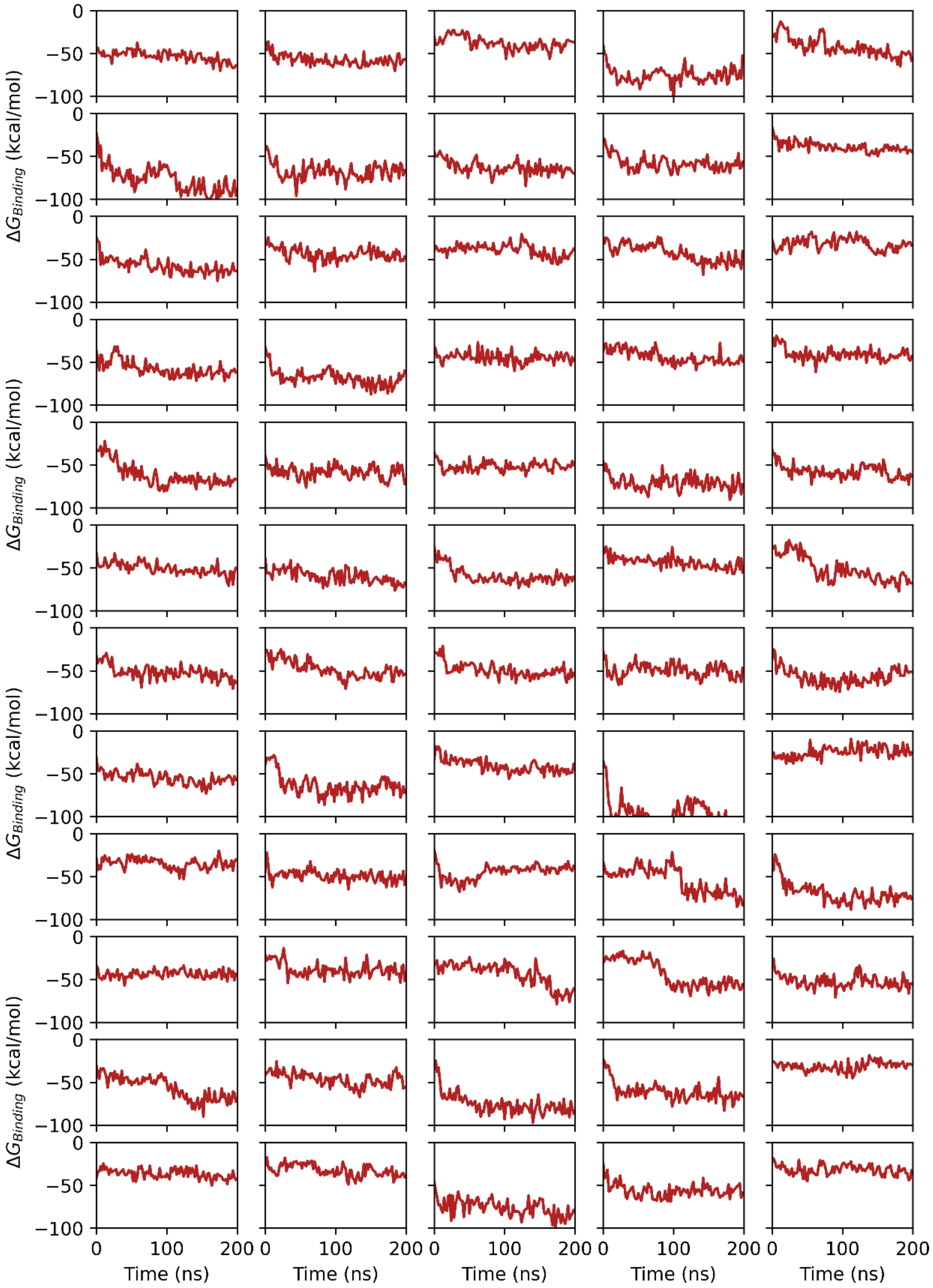


**Supplementary Figure 25. (b)** Results from samplings 61-120 are displayed.

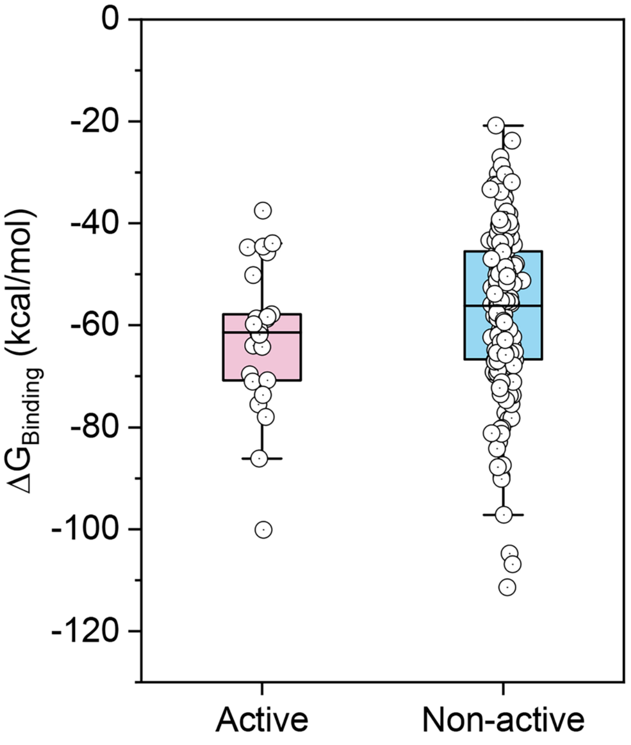


**Supplementary Figure 25. (c)** Distribution of binding free energy for active and non-active binding states of HotPETase.

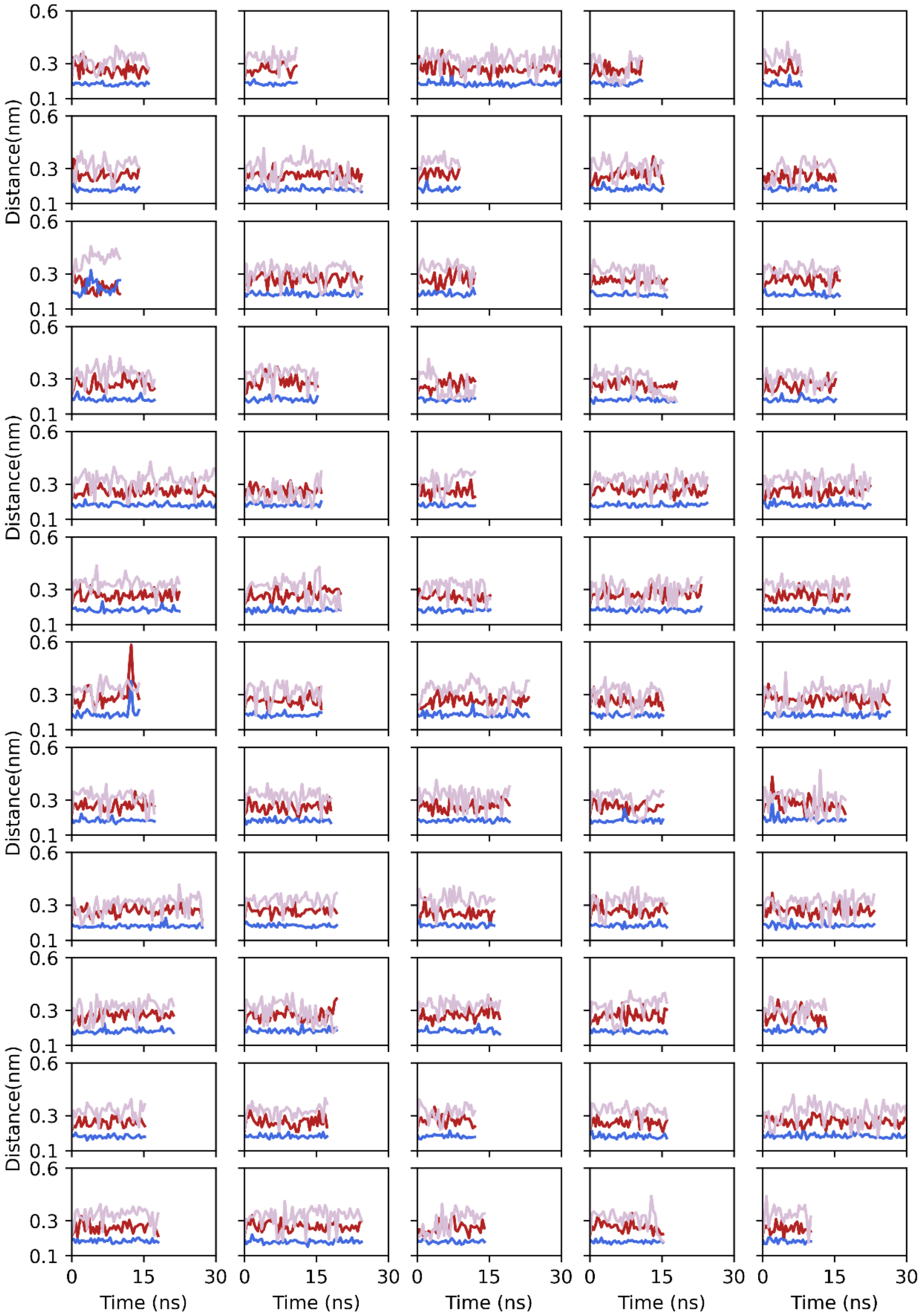


**Supplementary Figure 26.** (a) Three pairs of distances within the enzyme active site during supervised MD simulations of HotPETase. Each subfigure depicts the evolution of distances during one sampling. Results from samplings 1-60 are displayed.

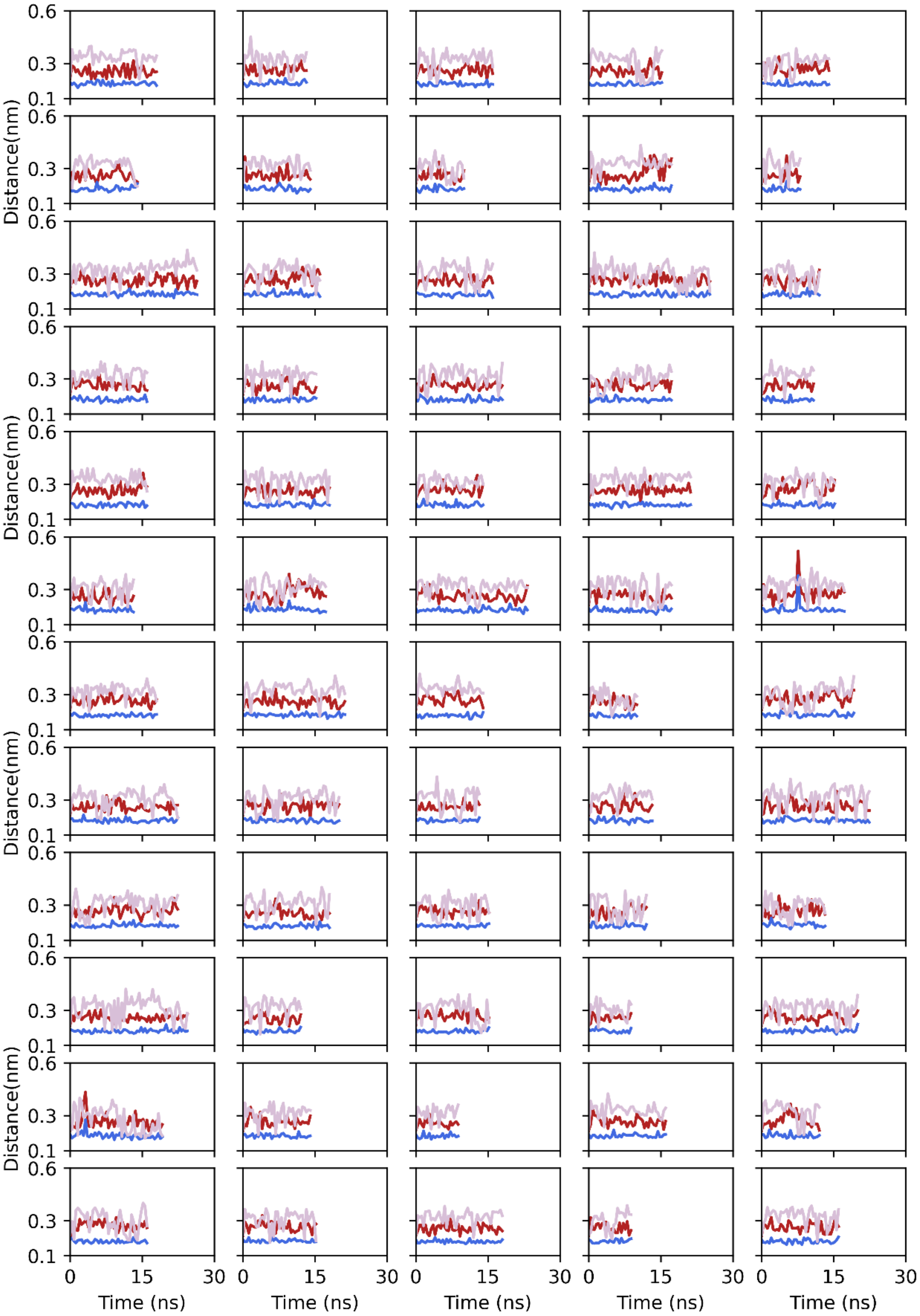


**Supplementary Figure 26.** (b) Results from samplings 61-120 are displayed.

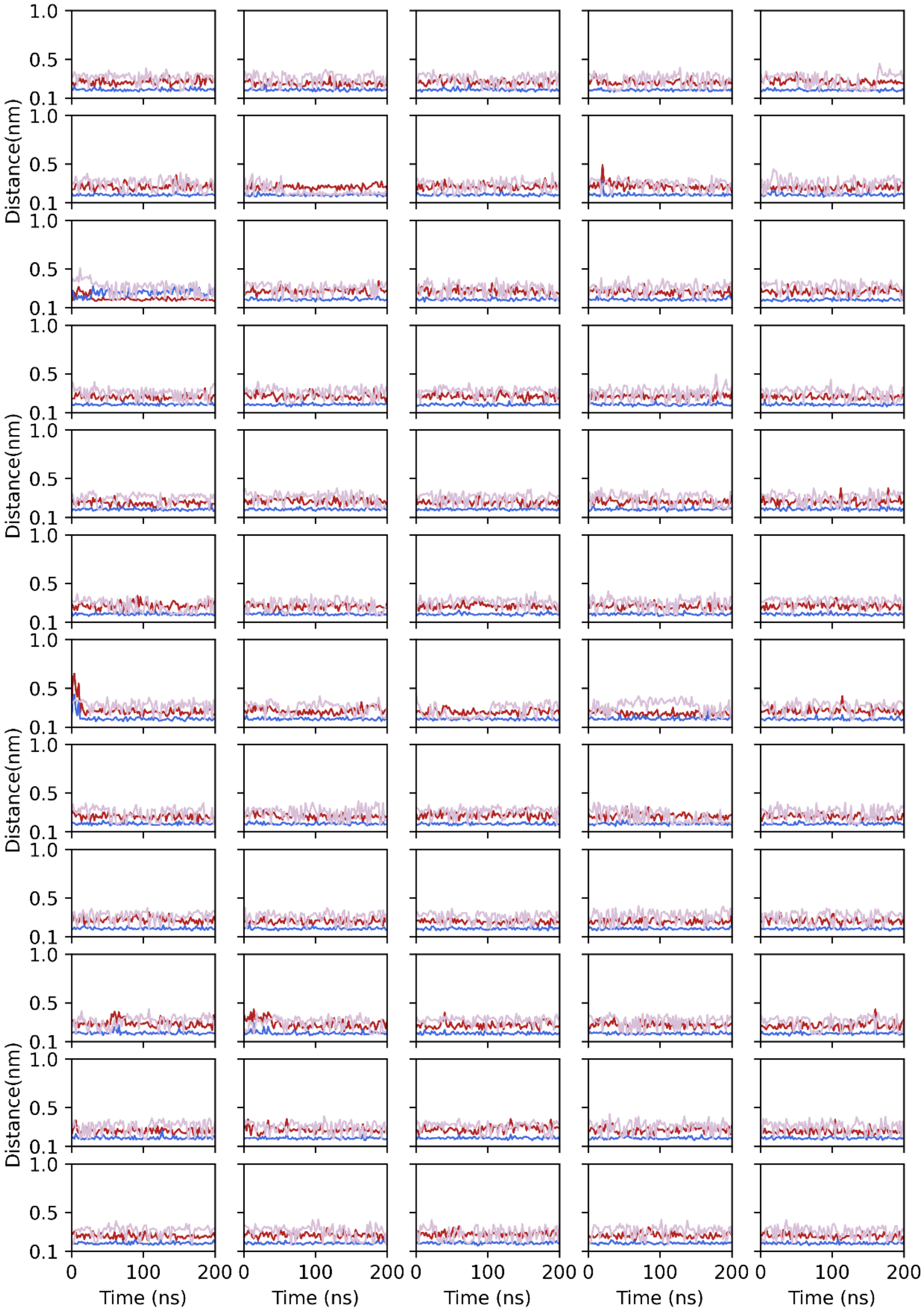


**Supplementary Figure 27.** (a) Three pairs of distances within the enzyme active site during 200 ns extended MD simulations of HotPETase. Results from samplings 1-60 are displayed.

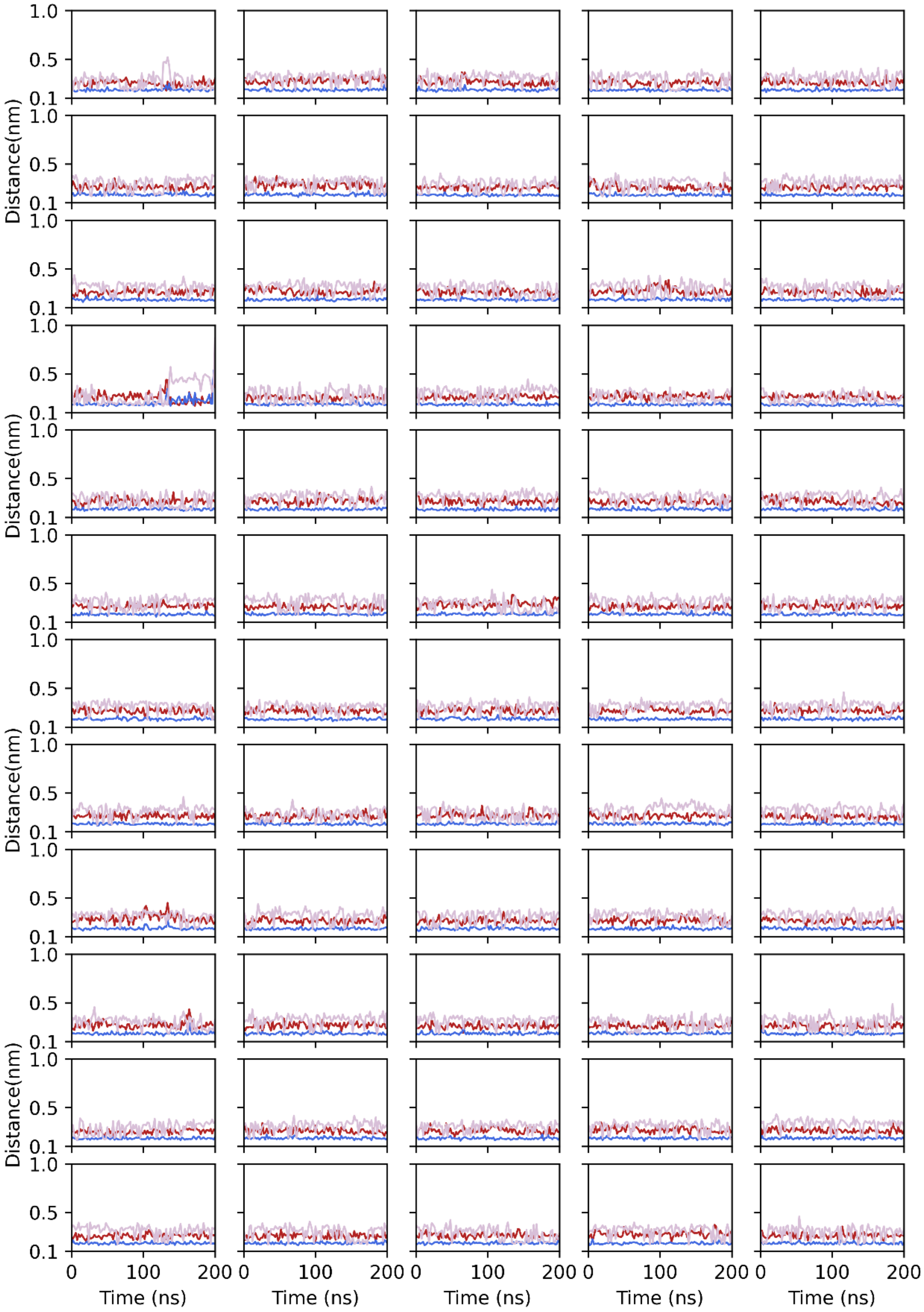


**Supplementary Figure 27.** (b) Results from samplings 1-60 are displayed.

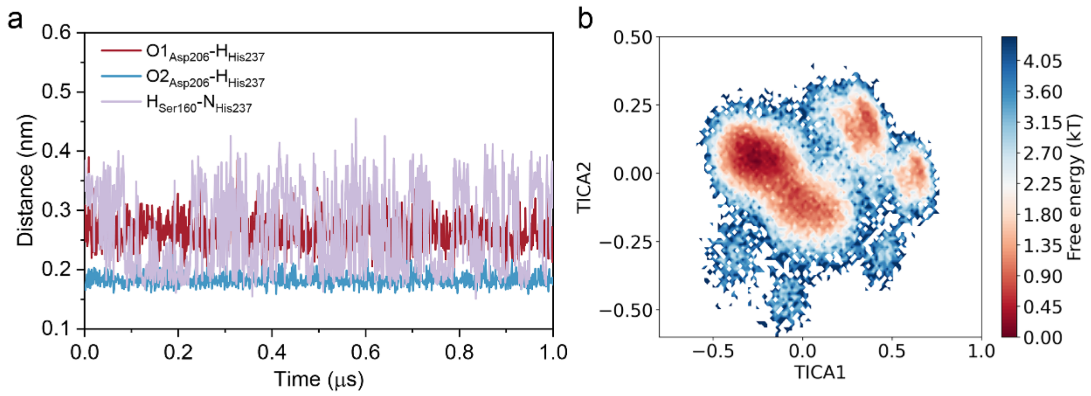


**Supplementary Figure 28.** (a) Three pairs of distances within the enzyme active site during 1μs-long MD simulation of unbound HotPETase. (b)The free energy landscape of unbound HotPETase was constructed using the TICA method for dimensionality reduction. The features utilized for this analysis included the side-chain torsion angles of ten residues in Supplementary Figure 18.

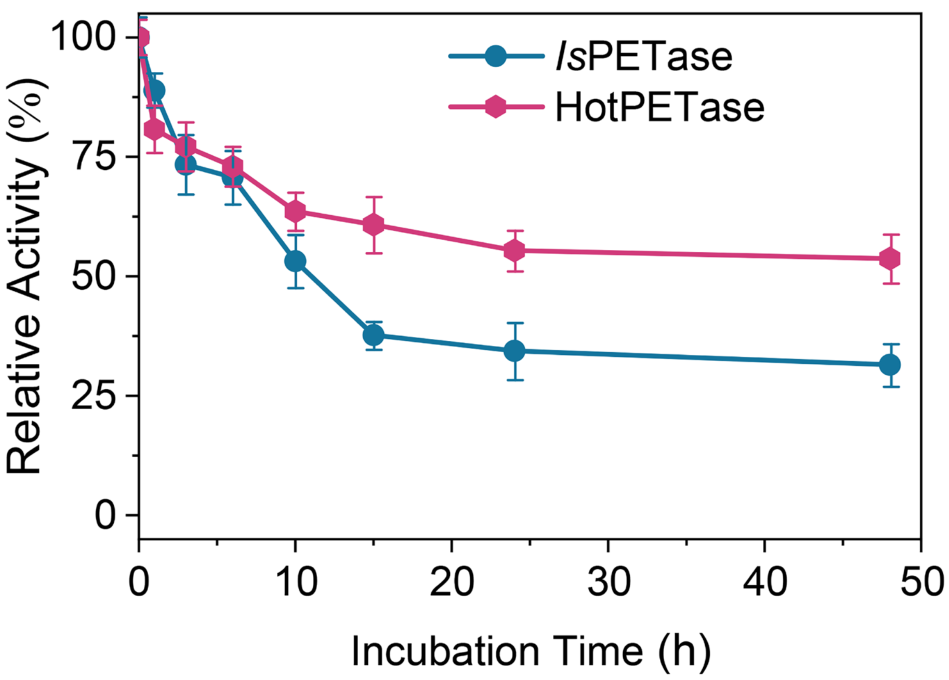


**Supplementary Figure 29.** Time-course deactivation of *Is*PETase and HotPETase without PET powder.

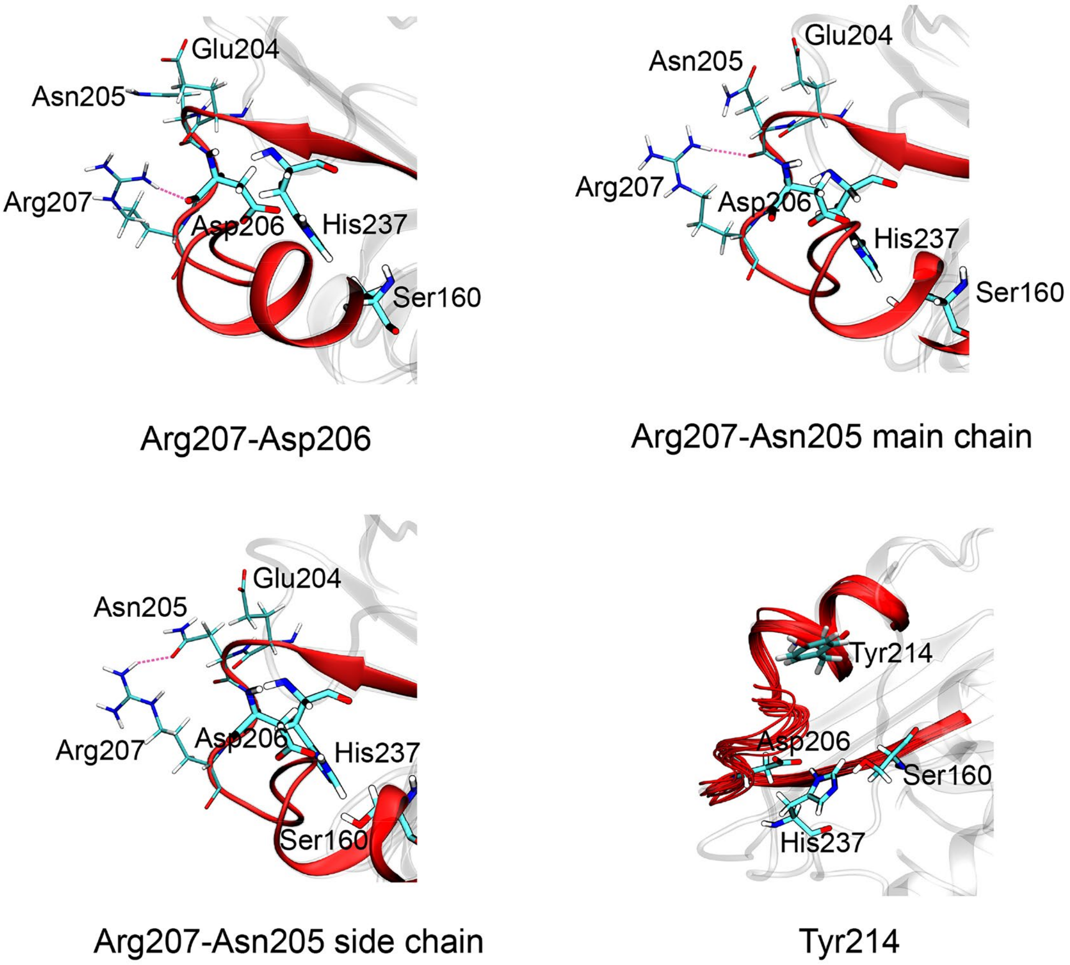


**Supplementary Figure 30.** The hydrogen bonds formed between Arg207 and other residues located in the catalytic loop (red) were illustrated. Additionally, the steric hindrance caused by Tyr214 was also depicted.

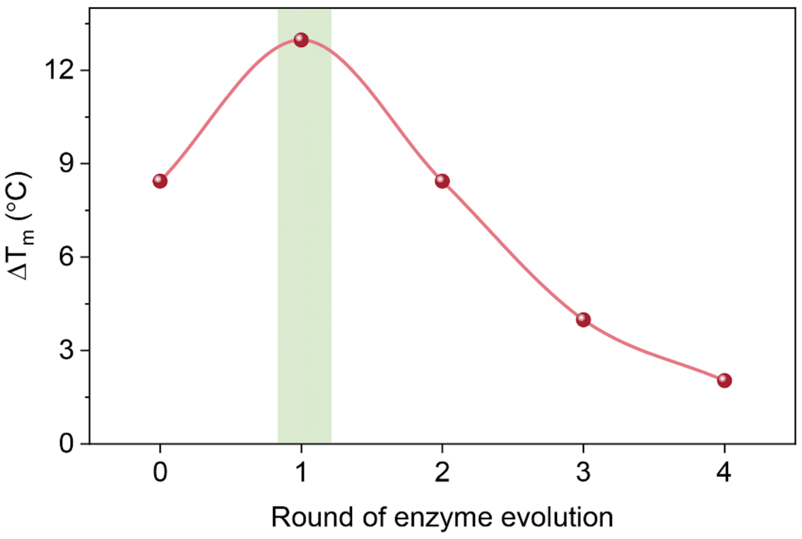


**Supplementary Figure 31.** The increase in the $T_m$ value of *Is*PETase variants during enzyme evolution. The thermostabilized variant of *Is*PETase, containing three mutations (S121E, D186H, R280A), referred to as *Is*PETase$^{TS}$, served as the starting point for engineering (round 0). Rounds 1-4 represent the protein variants obtained from the directed evolution of *Is*PETase$^{TS}$. The $T_m$ data extracted from Bell et al[3].

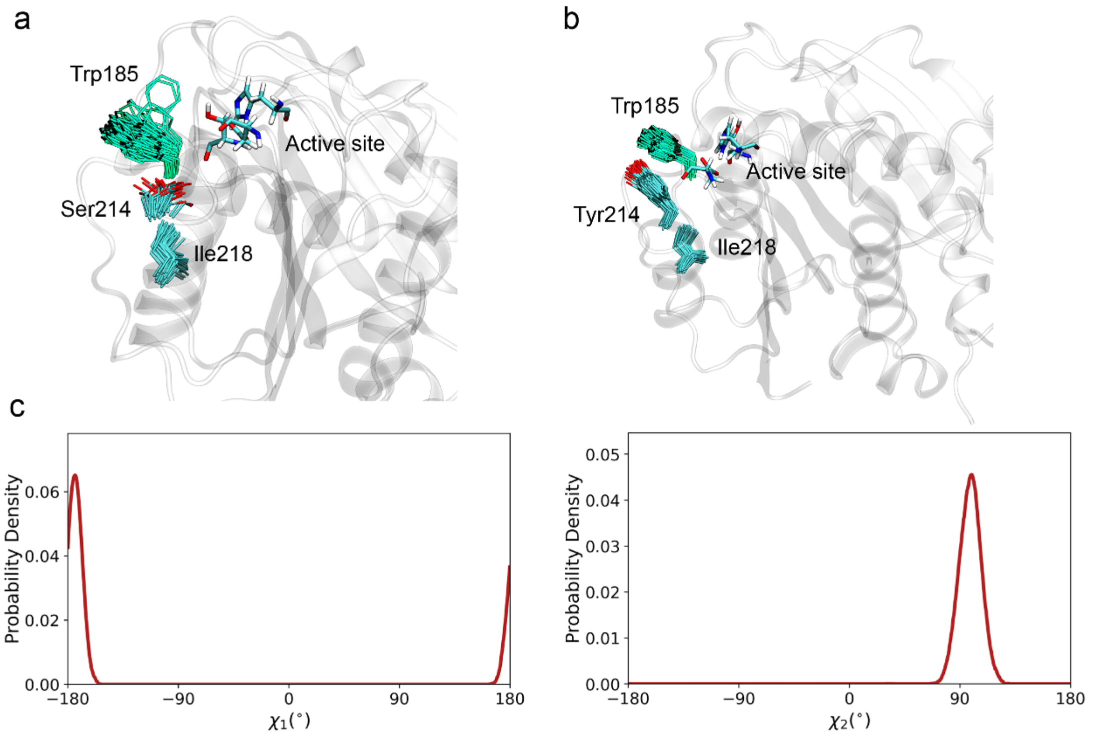


**Supplementary Figure 32.** Superimposed conformational states of Trp185, Ser214 and Ile218 in (a) *Is*PETase and (b) HotPETase. (c) Probability distribution of $\chi_1$ and $\chi_2$ of Trp185 in HotPETase.

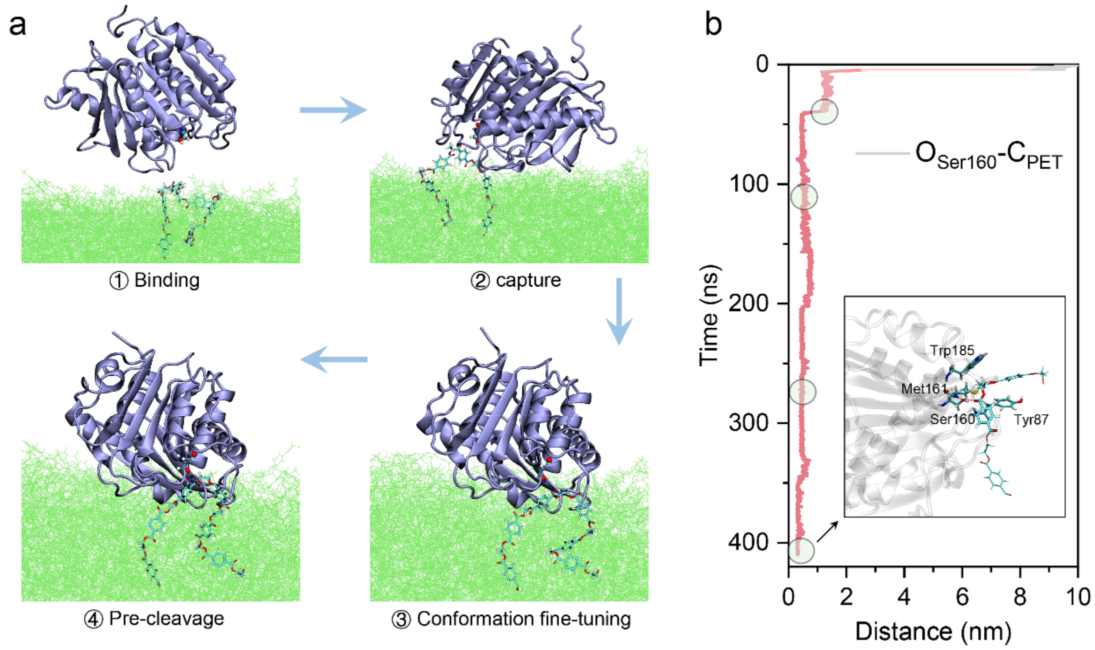


**Supplementary Figure 33. Ester bond recognition by HotPETase.** (a) Detailed mechanism of ester bond recognition by HotPETase. (b) Distance between the oxygen atom of the side chain hydroxyl of Ser160 and the carbon atom of the recognized PET ester bond.

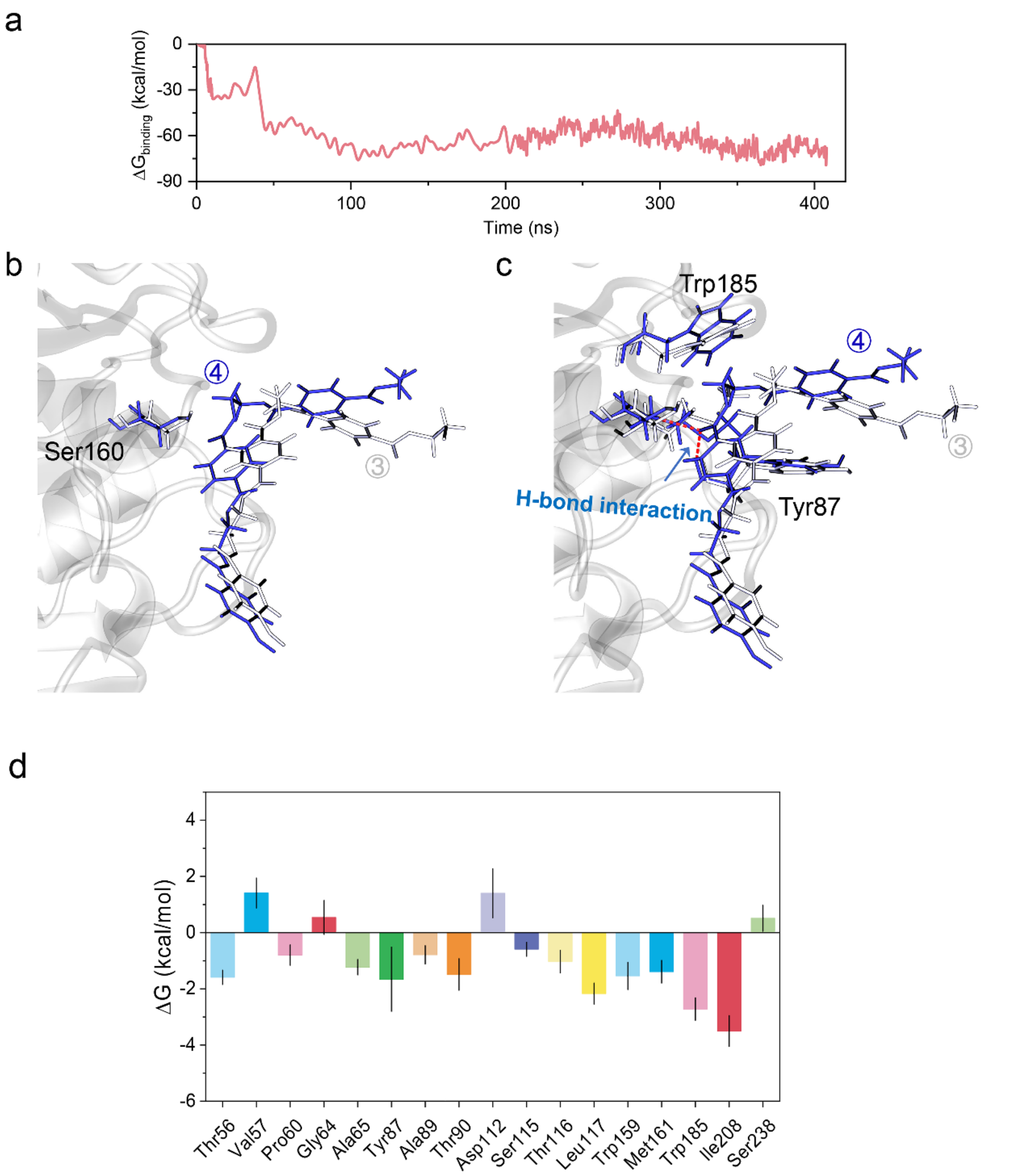


**Supplementary Figure 34.** (a) Binding free energy of IsPETase during supervised and extended (400 ns) molecular dynamics (MD) simulations. (b) The captured PET fragment exhibited close proximity after refinement of the binding conformation (③→④). The ③ and ④ represent the conformational refinement and pre-cleavage states. (c) Following the conformational refinement (③→④), the PET fragment formed an H-bond interaction with Tyr87 and Met161. (d) The top 17 residues of HotPETase contributing the most to the binding free energy with PET fragments.

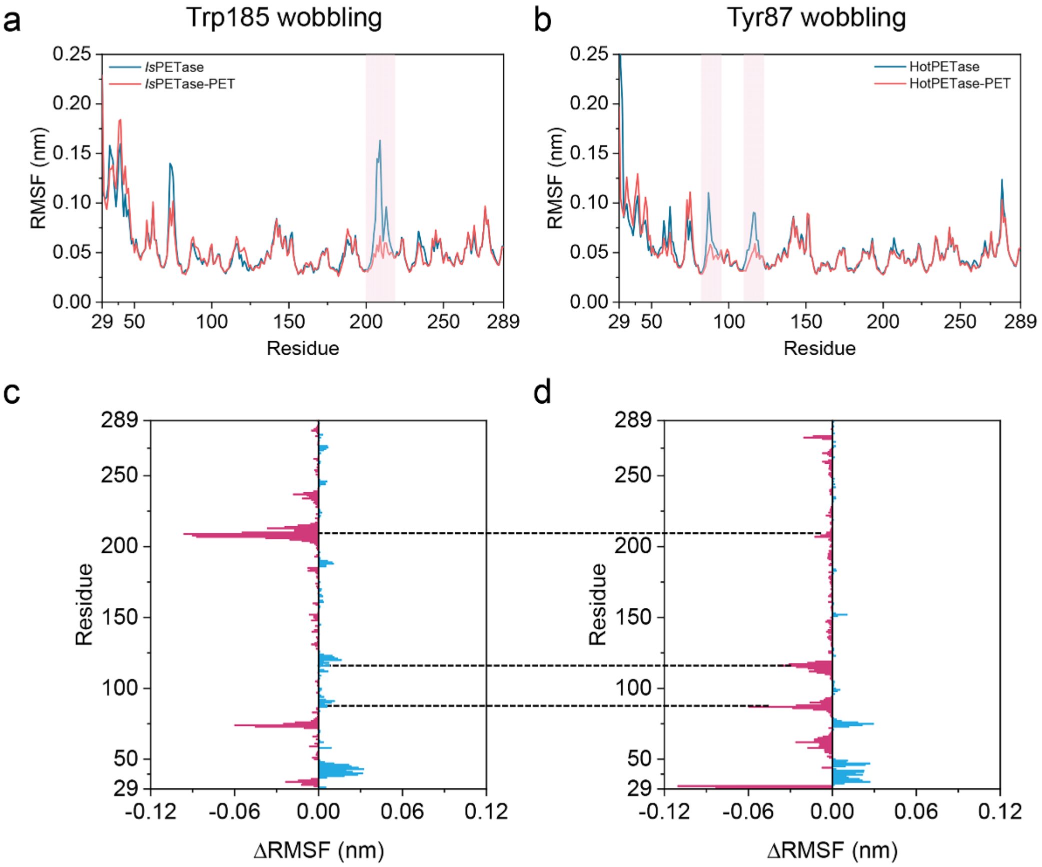


**Supplementary Figure 35.** The RMSF of bound and unbound states of (a) *Is*PETase and (b) HotPETase. The RMSF difference from unbound to bound was also shown (c for *Is*PETase, and d for HotPETase). In *Is*PETase, the RMSF of the catalytic loop (Asp206 located) showed a significant decrease in the PET-bound state. Additionally, the wobbling of Trp185 was also suppressed in the PET-bound state (Supplementary Figure 28). In HotPETase, the fluctuation of the catalytic loop was markedly reduced. The loop region containing Tyr87 exhibited higher fluctuation than that in *Is*PETase, which was subsequently reduced in the PET-bound state. We refer to this phenomenon as "Tyr87 wobbling".

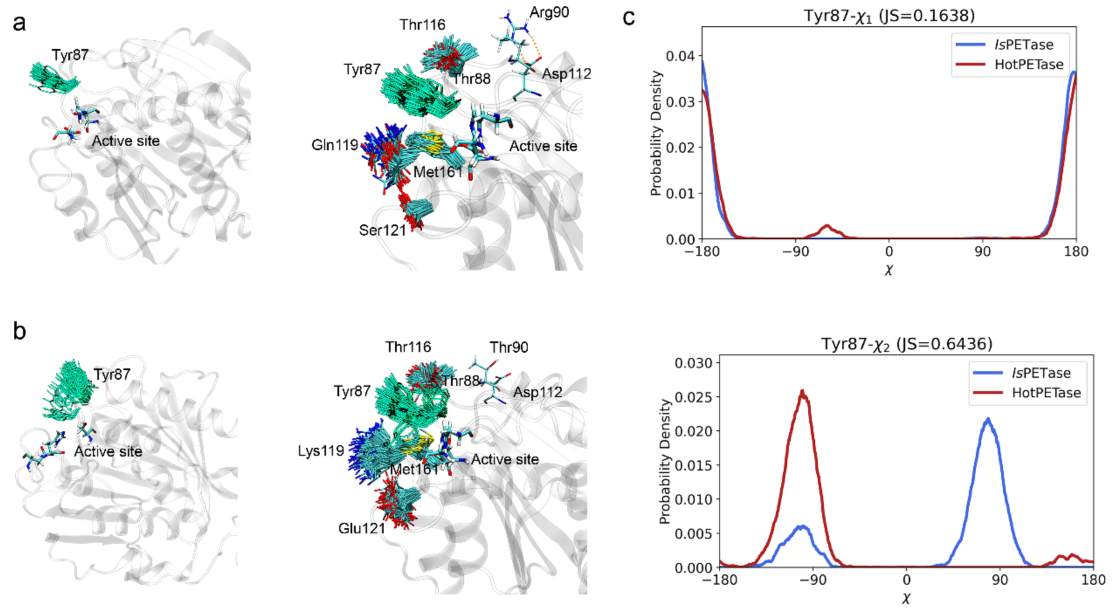


**Supplementary Figure 36.** Superimposed conformational states of Tyr87 and several related residues in (a) *Is*PETase, and (b) HotPETase. (c) Probability distribution of $\chi_1$ and $\chi_2$ of Tyr87 in *Is*PETase and HotPETase.

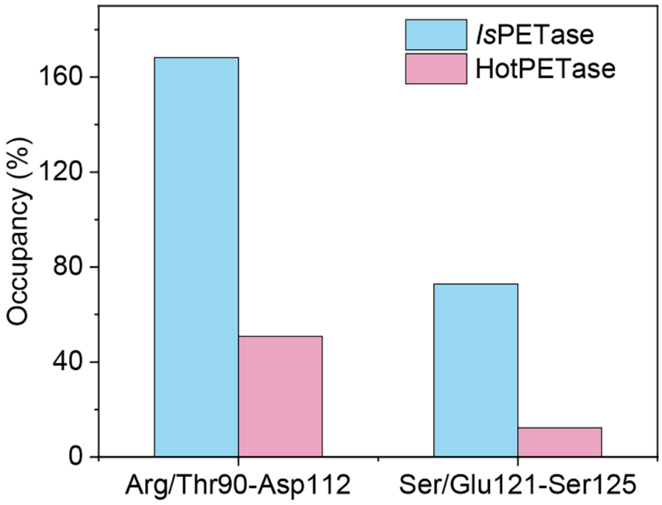


**Supplementary Figure 37.** Disruption of salt bridges or hydrogen bonds in HotPETase compared to *Is*PETase. When the occupancy exceeds 100%, it indicates that more than one hydrogen bond was formed between two residues. In *Is*PETase, a stable salt bridge exists between Arg90 and Asp112, whereas in HotPETase, there is a weaker hydrogen bond between Thr90 and Asp112. Additionally, a relatively stable hydrogen bond is formed between Ser121 and Ser125 in *Is*PETase, but this bond is disrupted in HotPETase due to the mutation of Ser121 to Glu. These two mutations result in a flexible conformation of the loop containing Tyr87 in HotPETase, leading to "Tyr87 wobbling."

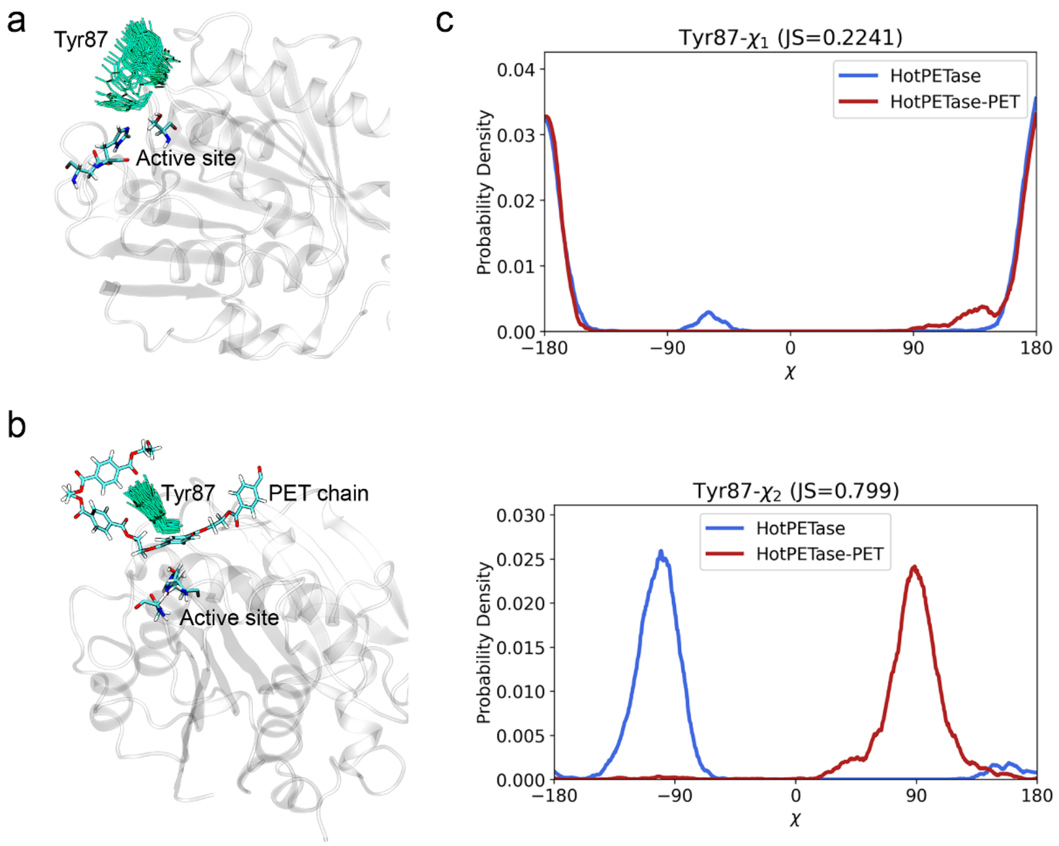


**Supplementary Figure 38.** Conformational difference of Tyr87 between (a) unbound and (b) PET-bound HotPETase. (c) Probability distribution of $\chi_1$ and $\chi_2$ torsion angles of Tyr87 in HotPETase and PET-bound HotPETase.

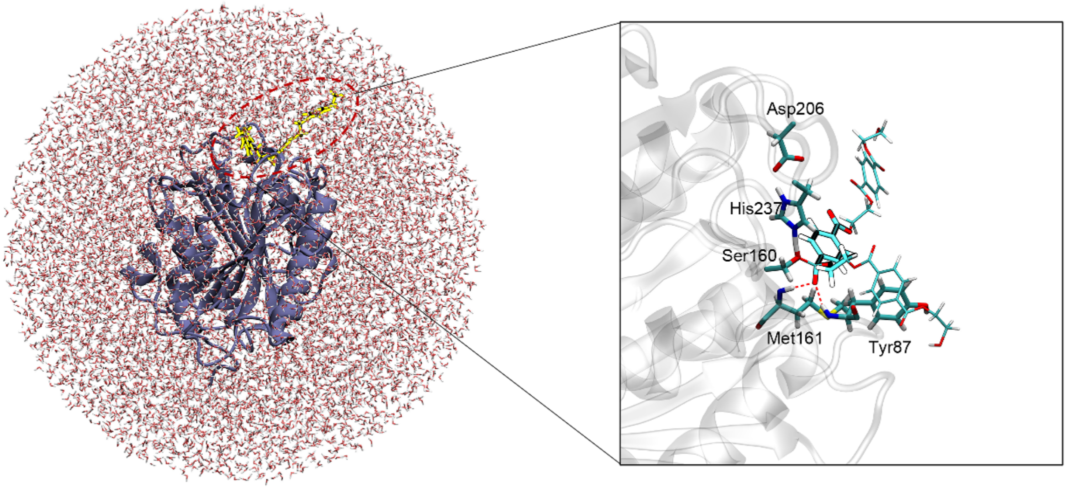


**Supplementary Figure 39.** QM/MM model in our simulations.

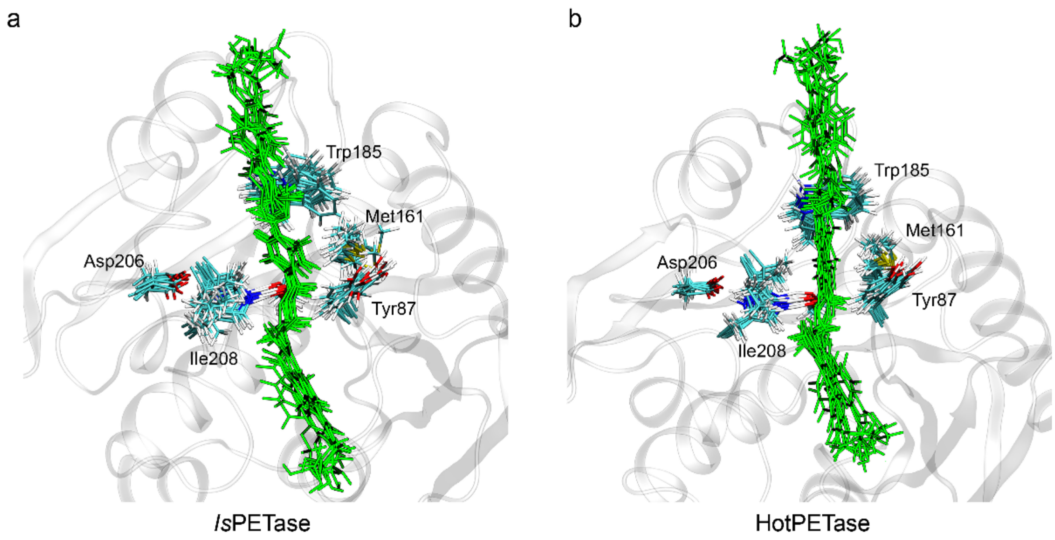


**Supplementary Figure 40.** Transition states of (a) *Is*PETase and (b) HotPETase.

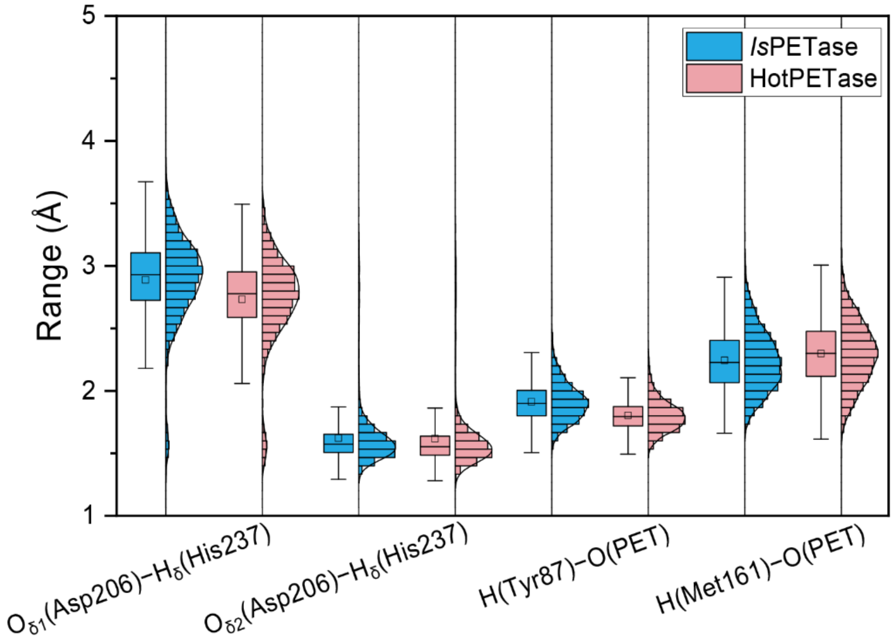


**Supplementary Figure 41.** A comparison of interatomic distances for four atom pairs within the transition states of *Is*PETase and HotPETase.

**Supplementary Table 1.** The enthalpic and entropic contribution of $\Delta G^{\ddagger}$ of *Is*PETase and HotPETase.

| Enzyme | $\Delta H^{\ddagger}$ (kcal/mol) | $\Delta S^{\ddagger}$ (kcal/mol/K) | $T\Delta S^{\ddagger}$ (298.15 K, kcal/mol) | $\Delta G^{\ddagger}$ (298.15 K, kcal/mol) |
|---|---|---|---|---|
| *Is*PETase | 16.3 | -0.008 | -2.4 | 18.7 |
| HotPETase | 13.4 | -0.013 | -3.9 | 17.3 |
